# Personality development in wild house mice: Evidence for a nutrition-dependent sensitive period early in life

**DOI:** 10.1101/2024.07.23.604785

**Authors:** Nicole Walasek, Milan Jovicic, Anja Guenther

**Author notes:** Correspondence concerning this article should be addressed to Nicole Walasek, Science Park 904, P.O. Box 94240, 1090 GE Amsterdam The Netherlands.

## Abstract

Changing environmental conditions pose serious challenges to organisms, for example, by disrupting access to food. Across species and traits, animals use phenotypic plasticity to rapidly adjust to such changes. Previous work has demonstrated that wild house mice are able to adjust stress coping to changing food quality within just three generations. However, we do not know when during ontogeny changing conditions induce phenotypic adjustments. We tested experimentally when during ontogeny (as fetus, newborn, weanling, or late adolescent) a food switch between standard and high-quality food shapes personality development (stress coping and stress perception) in cage-housed, wild house mice (*Mus musculus domesticus*). Personality traits were assessed in the Open Field and the Elevated Plus Maze at different time points during ontogeny (weaning, early adolescence, late adolescence, and adulthood). We observed three key findings. First, as mice grow older they tend to use more passive stress-coping strategies, indicating higher risk aversion. This relationship holds irrespective of food quality. However, mice fed with high-quality food show, on average, more active stress coping compared to mice receiving standard-quality food. Second, the fetal life stage might be a sensitive period for stress coping in response to experiencing decreases in nutritional quality. Third, experiencing an increase in nutritional quality may slow the age-related switch towards a passive stress-coping strategy. Our findings contrast previous work observing passive stress coping in mice living in semi-natural enclosures fed with high-quality food. We propose that the social environment of mice living in cages vs mice living in small groups may explain these differences. Our results highlight the need for experiments across the breadth of development comparing captive and semi-free-living animals. Ultimately, such studies will help us understand the complex relationships between development, nutrition, the (social) environment, and personality.

## Introduction

We live in a time of rapid environmental change. Although humans directly experience the impact of such changes (e.g. through extreme weather events), they are also indirectly affected through other species. For example, global environmental change (e.g. climate change, human-induced habitat modification, pollution) has led to declining biodiversity in insect populations (Wilson & Fox, 2021; Ziesche et al., 2023). This decline in insect biodiversity has a tremendous impact on crops; insects, which are natural predators of crop pests, are dying, while pests are thriving (Raven & Wagner, 2021). Human cities are populated by various rodent species that inevitably need to adapt their behavior to changes in climate. These adaptations have consequences for the spread of rodent-borne infectious diseases to which humans are susceptible (Beermann et al., 2023). Thus, understanding how non-human animals cope with changing environments can help us predict the consequences for humans.

One way in which organisms adapt to changing environments is through phenotypic plasticity (Agrawal, 2001; Charmantier et al., 2008). Within and across generations, plasticity allows organisms to adjust their phenotypes based on experiences during ontogeny (Bradshaw, 1965; DeWitt et al., 1998; Pigliucci, 2005; Taborsky, 2017; West-Eberhard, 2003; Whitman et al., 2009). These adjustments operate on faster time scales than natural selection and are often preceding genetic changes (Levis & Pfennig, 2016; B. W. Perry et al., 2018; Pfennig et al., 2010). Examples of phenotypic plasticity in nature abound. One example of the dramatic impact of phenotypic plasticity is the ability of coral fish to change sex in response to shifts in the populations’ sex-ratio (Casas et al., 2016; A. N. Perry & Grober, 2003).

Among invertebrates, water fleas and freshwater snails develop morphological defenses in response to changes in predator density (Herzog et al., 2016; Hoverman & Relyea, 2007; Relyea, 2003; Tariel-Adam et al., 2023). Across mammals, individuals can adjust reproductive timing, litter size, as well as body mass and size, in response to resource fluctuations caused by climate change (Boutin & Lane, 2014; Canale et al., 2016). These examples illustrate how environmental change can affect different dimensions of an organism’s ecology, such as their social environment, the prevalence of predators, and the abundance of resources. Each dimension brings about unique challenges and requires unique adaptations.

Changes in food availability or nutritional quality can have drastic consequences for organisms’ development. Nutrition across ontogeny is an important determinant of both physiological and behavioral development: The quality, abundance, and timing of available food determines when and how much organisms can grow, shaping their mating and reproductive strategies (Buchanan et al., 2022; English & Uller, 2016; Krause et al., 2009, 2017; Krause & Naguib, 2014; Lindström, 1999; Monaghan, 2007). For example, in bulb mites (*Rhizoglyphus robini*), nutritional conditions during the final developmental stages strongly determine whether males mature as aggressive and strong fighters or benign and weak scramblers (Leigh & Smallegange, 2014; Smallegange, 2011). In consuming food, organisms not only experience physiological changes but they also learn about their environment (English et al., 2016). These changes in the physiological and information state can in turn elicit changes in behavior. Such state-behavior feedback loops play a role in the development of individual differences in behavior (Ehlman et al., 2022; Mathot et al., 2017).

Over the past decade, interest has grown in understanding how changes in nutritional conditions shape individual differences in behavior (Beyts et al., 2023; Han & Dingemanse, 2015, 2017; Moran et al., 2021; Royauté & Dochtermann, 2017). Among-individual differences that are consistent across time and contexts (statistically indicated by being repeatable), are typically referred to as ‘animal personality’ (Coppens et al., 2010; Réale et al., 2007; Wolf & McNamara, 2012). A recent meta-analysis suggests that poor nutritional conditions tend to be associated with greater risk-taking behavior – implying lower levels of risk aversion – across various species (Moran et al., 2021). For example, mustard leaf beetles (*Phaedon cochleariae*) reared on low-quality food were bolder as adults compared to beetles who received high-quality food (Tremmel & Müller, 2013). These changes in boldness likely serve the purpose of increasing foraging success. Nutritional conditions also shape sexual signaling. A large body of work in song birds shows, for example, that nutritional deprivation during development compromises song production (Buchanan et al., 2022; Dougherty, 2021). In contrast, nutritional enrichment appears to promote higher investment into sexual signaling in various species (Buchanan et al., 2022; Porwal et al., 2023). At present we still lack a clear understanding of how exactly nutrition shapes the development of animal personality (Langenhof & Komdeur, 2018; J. A. Stamps & Groothuis, 2010; J. Stamps & Groothuis, 2010).

A well-rounded approach to studying how nutrition shapes animal personality involves embracing the dynamic interplay between evolution, development, physiology, and behavior (Duckworth, 2010; Duckworth et al., 2018; McElreath et al., 2007; Réale et al., 2007; J. A. Stamps & Groothuis, 2010; Trillmich et al., 2018; Wolf et al., 2007; Wolf & McNamara, 2012). Studies of animal personality have since moved towards exploring correlations between life-history traits (e.g., reproduction and growth), physiological traits (e.g., hormonal and metabolic traits), and behavioral traits in response to different environmental conditions and selection pressures (Dammhahn et al., 2018; Guenther et al., 2018; Laskowski et al., 2021; Réale et al., 2010).

Here, we highlight two recent studies in wild house mice (*Mus musculus domesticus)* living in semi-natural conditions (Lopez-Hervas et al., 2024; Prabh et al., 2023). Both studies explored how changes in nutritional quality shape stress-coping behaviors and life-history traits. Prabh et al. (2023) showed that mice are able to adjust life-history traits and behaviors within just three generations of experiencing changes in nutrition. Mice that underwent a switch towards calory-dense, high-quality food showed higher fecundity and shorter generation time, as well as more passive (risk-averse) stress-coping compared to mice fed with standard-quality food. These differences became more pronounced across generations. In a subsequent study, Lopez-Hervas et al. (2024) contrasted trans- and within generational effects of changes in nutritional conditions in the same population of mice. The authors found that adult mice were able to flexibly adjust life-history traits (reproduction, weight, and growth rates but not survival) to changes in food quality. However, adult mice were not able to adjust stress-coping behaviors. Irrespective of food quality, offspring of parents who experienced a food switch, developed more active stress-coping strategies and slower growth rates compared to mice experiencing stable nutritional conditions. For both behavioral and life-history traits, individuals who experienced a ‘loss’ in nutritional quality showed stronger responses, implying condition-dependent adjustment.

Taken together, these studies suggest that adult mice adjust life-history, but not stress-coping traits in response to changing nutritional conditions. This implies that the window of plasticity for modulating stress-coping has closed before adulthood. Developmental windows during which experiences exert the largest effect on development are called ‘sensitive periods’ (Fawcett & Frankenhuis, 2015; Frankenhuis & Fraley, 2017; Frankenhuis & Walasek, 2020). We speak of ‘critical periods’ when there is no plasticity outside of these windows (Knudsen, 2004). It remains an open question when during ontogeny changes in nutritional conditions have the largest effect on stress coping in rodents.

The present study provides first insights into sensitive periods for stress-coping in response to changing nutritional conditions in wild house mice (*Mus musculus domesticus)*. Various factors are known to shape coping behaviours, such as parental effects, social context, stress, and nutrition – and, studies recognize that these factors can have an especially strong influence at early life stages (Langenhof & Komdeur, 2018). However, existing studies rarely rigorously compare different ontogenetic windows both for the exposure to a specific experience and for measuring its effect (J. Stamps & Groothuis, 2010). Here, we present a study which has these qualities.

We experimentally manipulated the timing of changes in food quality in 407 cage-housed mice and measured its effect on stress-coping behaviors across ontogeny. Stress coping was measured at four time points during ontogeny in an Open Field and an Elevated Plus Maze. Measurements, in these forced exploration tasks predict risk-taking tendencies in mice under ecologically relevant conditions (Krackow, 2003; Krebs et al., 2019). We collected data in ‘control mice’ (never experienced a food-switch) and in ‘treatment mice’ (experienced a food-switch at different times during ontogeny). Ultimately, we attempted to answer the following research questions: First, how does stress coping change as a function of food quality across ontogeny; second, when during ontogeny is stress-coping in mice most sensitive towards a food switch? We derive hypotheses for these research questions based on previous studies (Lopez-Hervas et al., 2024; Prabh et al., 2023).

Regarding the first research question, we expect mice fed with high-quality food to display a higher tendency for passive stress coping compared to mice fed standard-quality food. We expect the duration of diet exposure to be positively associated with the strength of that diet’s effect. Thus, we expect differences in stress-coping between the standard- and high-quality food group to be more pronounced in older individuals. We do not have prior expectations about whether food type moderates the relationship between age and stress coping.

Regarding the second research question, we do not expect a food switch in older mice, approaching adulthood, to modulate behaviour. However, we expect that a food switch earlier in ontogeny (in utero, at birth, or at weaning) modulates behavior. We do not have prior expectations about which of these ontogenetic time points exerts the strongest effect. As with the first research question, we expect that any differences resulting from a food switch should be more pronounced in older individuals.

For transparency, all our data and code, including a preregistration will be made available on the open science framework upon publication. At present, we provide a view-only link to the preregistration to maintain the anonymity of the authors ([preregistration](https://osf.io/v7drq/?view_only=b4eda60ec8394e38ba27940459b97b4c)).

## Methods

### Animals and housing

Mice used to breed experimental mice for the experiments reported here descended from four semi-naturally housed populations of wild house mice (*Mus musculus domesticus*), originally caught in the Cologne/Bonn region of Germany. In brief, semi-natural populations consist of 100-200 individuals each. Each two of the semi-natural populations received standard-quality (SQ) food (Altromin © 1324), containing a total of 3227 kcal/kg metabolizable energy (24% protein, 11% fat, 65% carbohydrates) and high-quality (HQ) food (Altromin © 1414), containing ∼12% more metabolizable energy, i.e., 3680 kcal/kg (28% protein, 22% fat, 50% carbohydrates).

We removed young mice of the fifth generation shortly after independence (and before they started to be reproductively active) from the enclosures and let them habituate to being housed in a cage for at least three months in same-sex pairs before establishing breeding pairs. Mice continued to receive the same food as in the previous generations (i.e., standard-quality or high-quality food). All mice used in this study are the direct descendants of these mice and were born and raised in cages. Breeding cages were 425×265×180mm to allow animals building species-typical nests, and housing cages were 425×265×150mm.

Cages were equipped with woodchips for bedding, nesting materials, two shelters, and various enrichment items (e.g., a running wheel, a seesaw or climbing materials) which were changed every 2-3 weeks. Light and temperature varied naturally but underfloor-heating prevented temperatures to fall below 10°C. Throughout the experiments, animals were weaned at four weeks of age. Thereafter, they were kept in same-sex family pairs. To allow individual recognition, all animals were marked with a subcutaneously implanted RFID chip.

### Data collection

The data have been collected for five minutes according to standard protocols used in the Open Field test and Elevated Plus Maze test (details are provided below). Videos were recorded and analyzed using Ethovision (Noldus) software.

Data were always collected in the morning between 6am-10am. Only one individual per cage was tested per day and each animal only conducted one test per day to avoid any carry-over effects. Cages were transported individually to the testing room immediately prior to the test. All mice were handled by tunnel-handling to reduce stress of capture prior to the test. The test setups were cleaned with 70% ethanol between trials. Light conditions during the test were constant at 200 lux. All animals were handled only by experienced experimenters.

### Behavioral tests

Stress-coping and stress perception were measured in the Open Field and Elevated Plus Maze. The Open Field consist of a 60 x 60 cm brightly illuminated arena, which quantifies the time spent in the central area (in %) and total distance covered (in cm). The former indicates how stressful this environment was perceived, with shorter durations indicating lower stress levels. The latter indicates how mice cope with this stress-inducing experience. Specifically, larger distances indicate active (or proactive) coping, while shorter distances indicate passive (or reactive) coping.

Compared to the Open Field, the Elevated Plus Maze, which consists of a cross-shaped maze with two open, bright arms and two enclosed, dark arms, is less stressful. The two bright arms oppose each other and so do the two dark arms. For the Elevated Plus Maze, we recorded the following behaviors: time spent in dark versus bright arms (in %) and the total distance covered (in cm). As with the Open Field, the proportion of time spent in the open arms of the maze (excluding the center area between arms) indicates how stressful this environment was perceived. Shorter amounts of time spent in this area indicate that the environment was perceived as less stressful. As in the Open Field, the distance covered in the maze is indicative of the stress-coping strategy with larger distances indicating active (or proactive) coping.

### Experimental design

The experimental design comprises four individual experiments, each introducing a food switch at a different time point during ontogeny (appendix Figure 1A). Across the four experiments mice experienced a food switch during the fetal stage (mother’s third trimester; experiment 1), at birth (experiment 2), at weaning (experiment 3), or in late adolescence (corresponding roughly to early adulthood; experiment 4). Once a food switch occurred, mice remained on this diet for the remainder of the experiment. Mice could be switched from standard to high-quality food and vice versa. For comparative purposes, we maintained two control groups of mice who have been continuously fed standard or high-quality food and did not experience a food switch. Figure 1 illustrates when in ontogeny mice were tested in the Open Field and the Elevated Plus Maze for each experiment (each mouse underwent both tests repeatedly; the testing order was Open Field first and Elevated Plus Maze at least 48 hours later).

**Figure 1:**
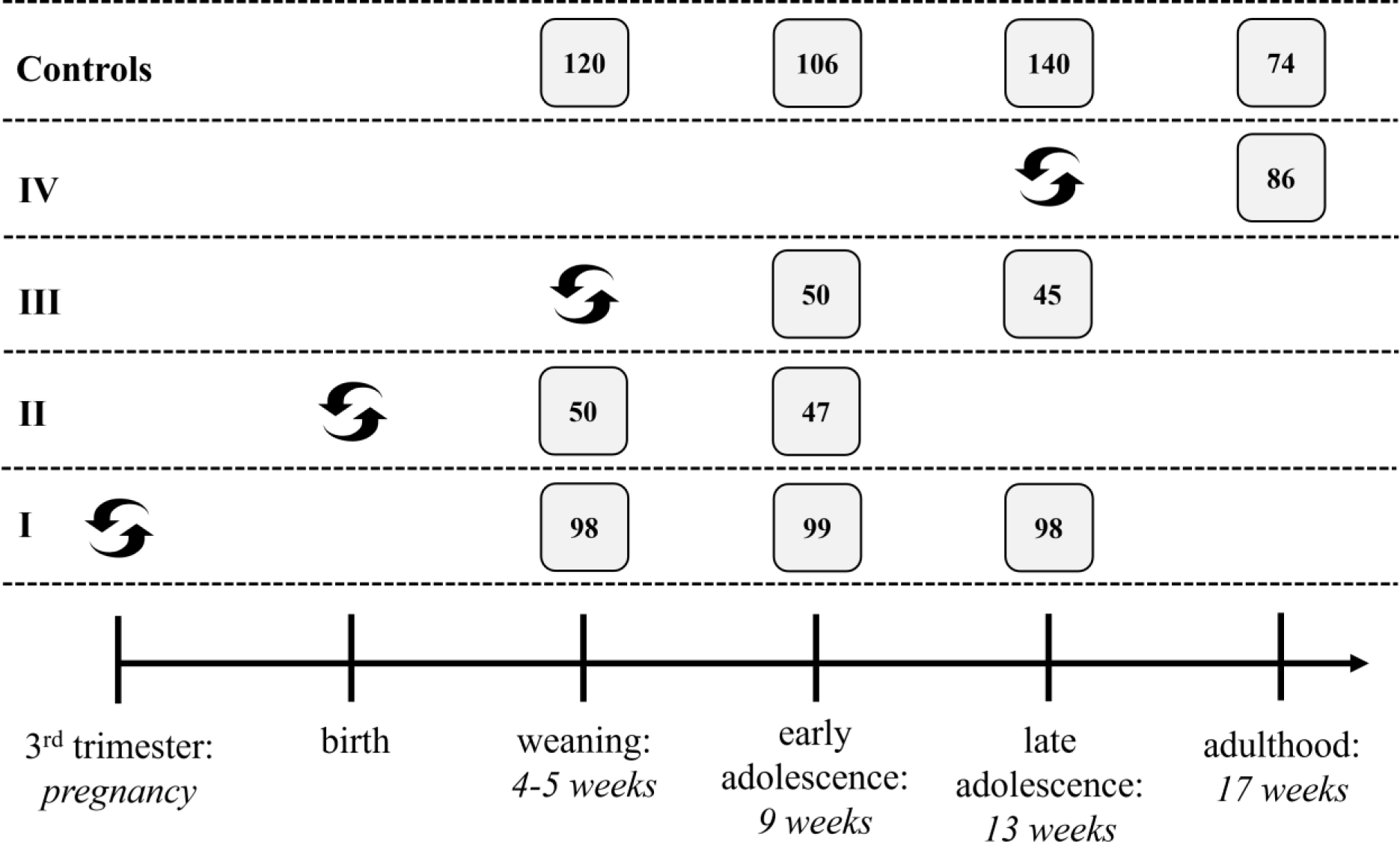
Overview over the food switch and measurement times across experiments. This figure illustrates when mice within each experiment were tested. The numbers within each square indicate the sample size of mice that underwent a specific treatment (indicated by the row) and were measured at a specific time (indicated by column). We report the sample sizes from the Open Field experiment. The sample size for control individuals fluctuates. The reason for this is that we replaced mice in different experiments such that control mice are not tested more often than treatment mice. Timing of food switch is indicated by the black, curved arrows and measurement times are indicated by grey squares. The horizontal axis indicates ontogeny, from the fetal stage to adulthood. Experiments are separated by rows and the roman numeric on the left indicates to which experiment the row belongs to. The top row shows measurement times in control-mice.

### Data set and variables

The final data set consists of 406 cage-housed mice with a total of 995 measurements from the Open Field and 973 from the Elevated Plus Maze (EP). The measures were taken across four different test ages (i.e., weaning, early adolescence, late adolescence, and adulthood). Of those 995 (EP: 973) measures, 440 (EP: 434) are from control individuals who never experienced a food switch and 555 (EP: 539) measures are from the treatment group who experienced a food switch. These numbers already account for missing data (see below). The dataset is both cross-sectional and longitudinal. It consists of four cross-sectional experiments: one for each food switch time. Within experiments mice have been assessed repeatedly between 1 – 3 times at different test ages (see Figure 1).

Aside from information about the food type, timing of food switch, and test age, additional variables indicate the sex of the mouse, mother ID, and cage ID.

### Missing data and outliers

The data contain 12 cases with missing values on cage ID. For these cages, we assign a unique cage ID. There are 2 cases with missing values on the Open Field outcome measures which will be excluded from the corresponding analyses using these outcomes. There are 22 cases with missing values on the Elevated Plus Maze outcome measures which were excluded from the corresponding analyses. None of the other variables contained missing values.

We did not exclude any statistical outliers except for one analysis. When analyzing distance covered in the Open Field of adult mice who were switched from high to standard-quality food, we removed three statistical outliers to improve model fit. Additionally, we excluded cases when the tracking software was unable to find the mouse for more than 20% of the time in the Elevated Plus Maze (N=8 cases). In contrast, mice could always be found in the Open Field, thus we retained all measures there.

### Statistical analyses

All analyses were performed in R (version 4.3.0; R Core Team, 2023). We computed repeatabilities of all four behavioral measures adjusted for age using the *rptR* package (Stoffel et al., 2017). To account for the nesting and repeated measures structure of our data, we fitted mixed-effects models from the *lme4* package (Bates et al., 2015). We assumed Gaussian distributions of outcomes in all models. We used the *performance* (Lüdecke et al., 2021) package to visually check assumptions and model fit (i.e., normality of residuals, linearity, and homogeneity of variance). The variable indicating time spent in the center of the Open Field (Center OF) was square-root transformed to approximate a Gaussian distribution and improve model fit. We used the packages *lmerTest* (Kuznetsova et al., 2017) and *pbkrtest* (Halekoh & Højsgaard, 2014) to approximate Kenward-Roger degrees of freedom for statistical inference (i.e., p-values and confidence intervals). Additionally, we computed bootstrapped p-values and confidence intervals using the package *parameters* (Lüdecke et al., 2020) to verify the robustness of our statistical inference. We find that all estimation methods lead to similar results. To control the false-discovery rate (i.e., significant findings that are false positives due to a large numbers of tests), we use the Benjamin-Hochberg procedure to adjust p-values across analyses. For each outcome measure of interest (distance in the Open Field, time spent in center of the Open Field, distance in the Elevated Plus Maze, and time spent in the bright arms of the maze), we ran separate models. Each of these models has the same set of predictors and only differs in said outcome. The p-value correction was applied across these models with different outcome measures. Thus, for each p-value of each predictor, we report a ‘corrected’ p-value, adjusted across the four outcome measures.

To estimate the effects of categorical predictors, we conduct analyses of variance (ANOVAs) with Kenward-Roger degrees of freedom based on the mixed models. We estimate type III sums of squares. When our research question requires it, we use the package *emmeans* (Lenth, 2023) to perform post-hoc comparisons between groups and to estimate marginal means. We apply the Tukey procedure to adjust for multiple testing during post-hoc comparisons.

We use two-tailed tests for all analyses with an alpha-level of 0.05 and 95% confidence intervals. Below we detail the structure of our models answering different research questions.

#### RQ1: How does behavior change as a function of food quality across development?

This analysis only uses mice from the control group. To answer the research question, we perform a moderation analysis with food type as the moderator for each behavioral outcome. We test whether food type moderates the relationship between test age and the outcome, while controlling for sex. In these models, we use nested random intercepts to account for the nesting of mice in cages and cages in families (i.e., random effect structure: family/cage/mouse). Due to their involvement in interactions, we use sum-to-zero coding for food type and test age. Here, and in all other analyses we use dummy coding for sex with females as the reference group. To test for differences between groups we conduct post-hoc comparisons.

#### RQ2: When during development are mice sensitive towards a food switch?

In order to ensure that we have a proper control condition for each treatment, we split the analysis into two sets of models: One set compares ‘control-mice’ who have been continuously fed standard-quality food to ‘treatment-mice’ who were switched to high-quality. The other set compares mice who have been continuously fed high-quality food and switched to standard food. Additionally, we run separate models for each test age (because possible adjustment times after a food-switch differ across test ages) and for each behavioral outcome measure. The general structure of each individual model is the following: We regress the outcome measure on switch time, while controlling for sex. In these models, we use nested random intercepts to account for the nesting of cages in families (family/cage). Switch time is dummy coded with never as the reference group. Thus, we do not need additional post-hoc comparisons to compare groups.

#### Exploratory analysis: Does the timing of food switch shape changes in behavior across development?

In this analysis, we aim to estimate how behavior changes in developing mice who experienced a food switch. For that purpose, we use mice from different test ages (combined from different experiments) who experienced a food switch at the same times. Specifically, we combine mice from Experiments 1 and 2 to estimate changes in behavior from weaning to early adolescence (possible switch times: birth and pregnancy). Then, we combined mice from Experiments 1 and 3 to estimate changes from early to late adolescence (possible switch times: pregnancy and weaning). We split the analysis into two sets with separate models for switches from standard- and high-quality food (as in RQ2). Then, we performed a moderation analysis with switch time as the moderator for each outcome measure. We tested whether switch time moderates the relationship between test age and the behavioral outcome, while controlling for sex. In these models, we used nested random intercepts to account for the nesting of mice in cages and cages in families (family/cage/mouse). Switch time and test age were sum-to-zero coded due to their involvement in interaction terms. In cases in which we discovered a significant moderation effect, we additionally perform post-hoc comparisons for each level of the moderator.

### Preregistration

We preregistered all analyses carried out in this study as a secondary data preregistration on the open science framework (<u>preregistration</u>). The data have not been used in a previous study and were specifically collected for the present study. However, the data were already collected at the time of writing the preregistration which is why we chose the template for secondary data analysis. We report the following four minor deviations from our preregistration. First, contrary to the information in the preregistration, we used sum-to-zero coding for RQ2 and the exploratory analysis. Using dummy-coding, as suggested in the preregistration, is suboptimal due to the involvement of interaction terms. Second, we suggested an additional exploratory analysis relying on training a classifier to discriminate behavior between standard and high-quality fed mice. We had to discard this analysis because the associations between food type and behavior were not strong enough to successfully train such a classifier. Third, there are slight deviations between the number of measures reported in the preregistration and here. The numbers in the preregistration do not account for the removal of 8 cases in which the tracking software did not find the mice for at least 20% of the time. Lastly, as described earlier, we had to remove three outliers from one analysis to improve model fit. This removal was only based on information obtained from assumptions checks.

### Ethics

Housing of wild mice is approved and regularly controlled by the Veterinäramt Plön under licence: 1401-144/PLÖ-004697. All animals were handled, and procedures were carried out according to national guidelines. For animals tested in the Open Field and Elevated Plus Maze tests, experiments were performed under licences V244 – 31223/2019(62-5/19). Reduced sample sizes during experiments result from male mice becoming aggressive towards each other. In such cases, we separated males and both dropped out of the experiments to avoid any potential influences of isolation on behavioral outcomes. In case a male was injured during aggressive encounters, we treated its wound and closely observed its health status until the mouse was healthy again or until we decided to euthanize (N =2 cases).

## Results

### RQ1: How does behavior change as a function of food quality across development?

All behaviors were significantly repeatable across age (Table 1). Food type did not moderate the relationship between age and any of the behavioral outcomes (appendix Tables A2-A3). However, we did find that age significantly predicted all behaviors (distance Open Field: *F* (3,321.25) = 22.70, *p* < .001; center Open Field: *F*(3,325.11)= 6.18, *p* < 0.001; distance Elevated Plus Maze: *F*(3,304.74) = 4.27, *p* < 0.006; bright arm Elevated Plus Maze: *F*(3,326.18) = 4.96, *p* < 0.003; appendix Table A1; post-hoc comparisons and marginal means in appendix Tables A3-A4 and Figure A3). Time spent in the center of the Open Field and in the bright arms of the Elevated Plus Maze tended to increase with age, indicating that older mice perceived the tests as less stressful compared to younger mice. At the same time, older mice tended to cover less distance in both the Open Field and the Elevated Plus Maze, indicating an increasingly passive stress-coping strategy. Across all four behaviors, food type only significantly predicted distance covered in the Open Field (*F*(1,73.66) =17.25, *p* < .0001; appendix Table A1, Figure 2), comparable to previous findings in the same population. Contrary to previous findings in mice from semi-natural enclosures (Lopez-Hervas et al., 2024; Prabh et al., 2023), we found that cage-housed mice on a high-quality diet covered more distance in the Open Field compared to the standard-quality food group (Figure 1). This result implies that high-quality food promotes an active stress-coping strategy in caged-house mice. As we are particularly interested in the impact of food type on development, we decided to focus on distance covered in the Open Field as our main behavioral outcome of interest for the remainder of the results. Figure 2 summarizes how distance in the Open Field changes in relation to age and food type.

**Table 1.** Repeatability estimates in control mice.

|  | *R* | *SE* | 2.5% CI | 97.5% CI | *p* (LRT) | *p* (permut) |
| --- | --- | --- | --- | --- | --- | --- |
| **Distance (OF)** | 0.390 | 0.053 | 0.283 | 0.489 | **<0.001** | **0.001** |
| **SQ** | 0.690 | 0.053 | 0.570 | 0.778 | **<0.001** | **0.001** |
| **HQ** | 0.214 | 0.081 | 0.051 | 0.361 | **<0.001** | **0.015** |
| **Center (OF)** | 0.244 | 0.057 | 0.132 | 0.355 | **<0.001** | **0.001** |
| **SQ** | 0.109 | 0.072 | 0.000 | 0.262 | 0.082 | 0.074 |
| **HQ** | 0.242 | 0.080 | 0.088 | 0.400 | **<0.001** | **0.001** |
| **Distance (EP)** | 0.425 | 0.053 | 0.318 | 0.529 | **<0.001** | **0.001** |
| **SQ** | 0.418 | 0.080 | 0.244 | 0.553 | **<0.001** | **0.001** |
| **HQ** | 0.403 | 0.078 | 0.256 | 0.561 | **<0.001** | **0.001** |
| **Brightarm (EP)** | 0.129 | 0.055 | 0.020 | 0.234 | **0.004** | **0.006** |
| **SQ** | 0.100 | 0.073 | 0.000 | 0.261 | 0.055 | 0.075 |
| **HQ** | 0.201 | 0.080 | 0.041 | 0.367 | **0.008** | **0.009** |

Table 1: Repeatability adjusted for age. For each outcome we show repeatability irrespective of food type and separately depending on food type. We use the following abbreviations: OF = Open Field, EP= Elevated Plus Maze, SQ = Standard-quality food, and HQ = High-quality food.
| *R* | *SE* | 2.5% CI | 97.5% CI | *p* (LRT) | *p* (permut) |
| --- | --- | --- | --- | --- | --- |

**Figure 2:**
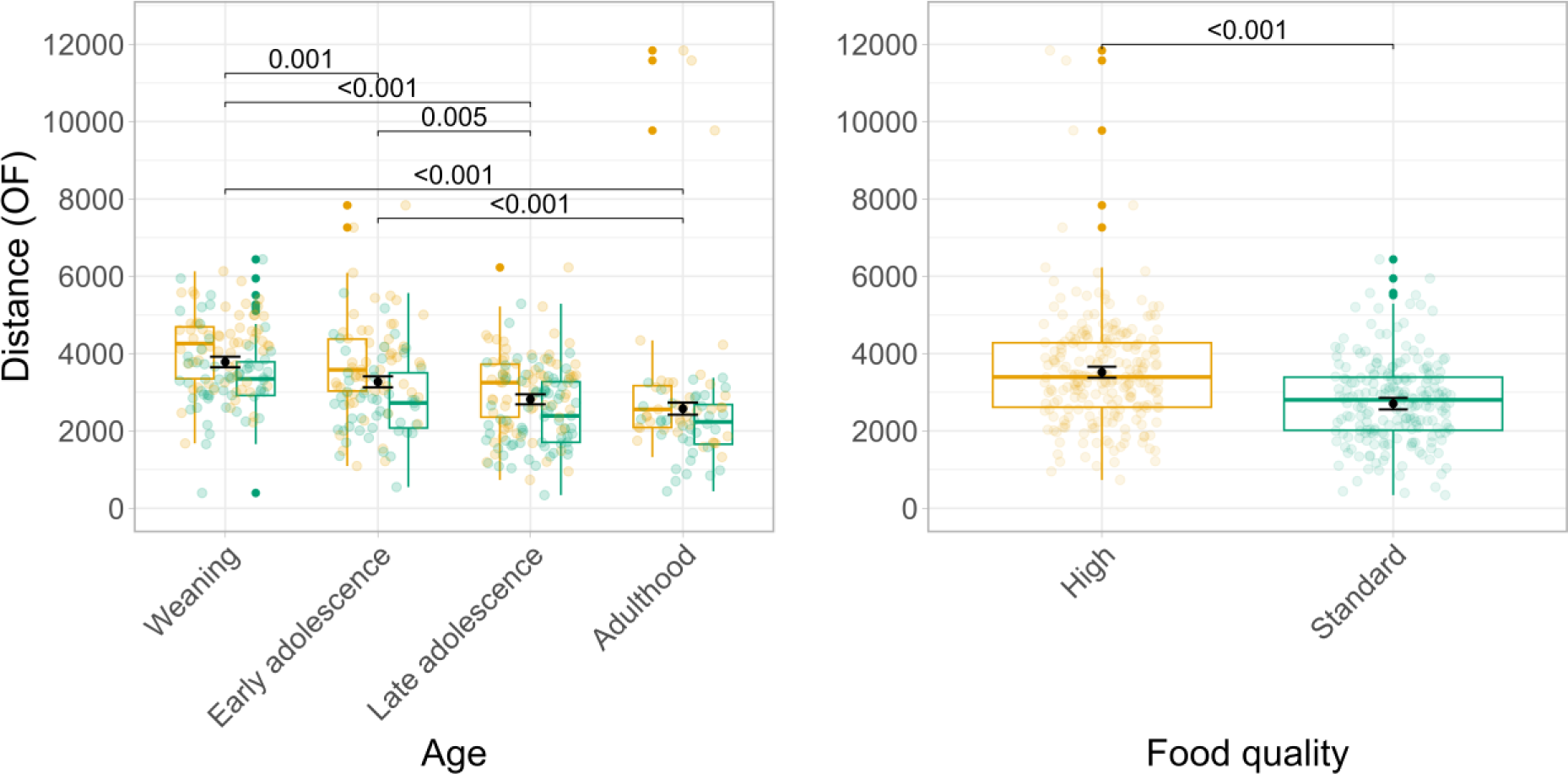
Distance covered in the Open Field across development. The left panel shows boxplots for distance covered in cm (y-axis) across test ages (x-axis). Colors indicate the food quality with yellow corresponding to high-quality and green to standard-quality food. The black points indicate model-based estimated marginal means for each test age and corresponding standard errors. P-values indicate significant differences between test ages.

The right panels illustrates the main effect of food quality (x-axis) on distance covered in the Open Field (y-axis). P-values indicate significant differences between food types. P-values have been corrected for multiple post-hoc comparisons with the Tukey procedure.

### RQ2: When during development are mice sensitive towards a change in food?

Our analysis revealed no significant effects on distance in the Open Field when switching mice from standard-quality to high-quality food at any of the tested switch times during ontogeny (Tables 3-4 and Figure 3 bottom row). An interesting descriptive observation is that the mean distance of mice whose diet was switched in utero was lower than the mean of mice who have been switched at birth or weaning. This is somewhat surprising as mice who experienced a food switch as fetuses, theoretically had the longest time to adapt to their new diet.

**Figure 3:**
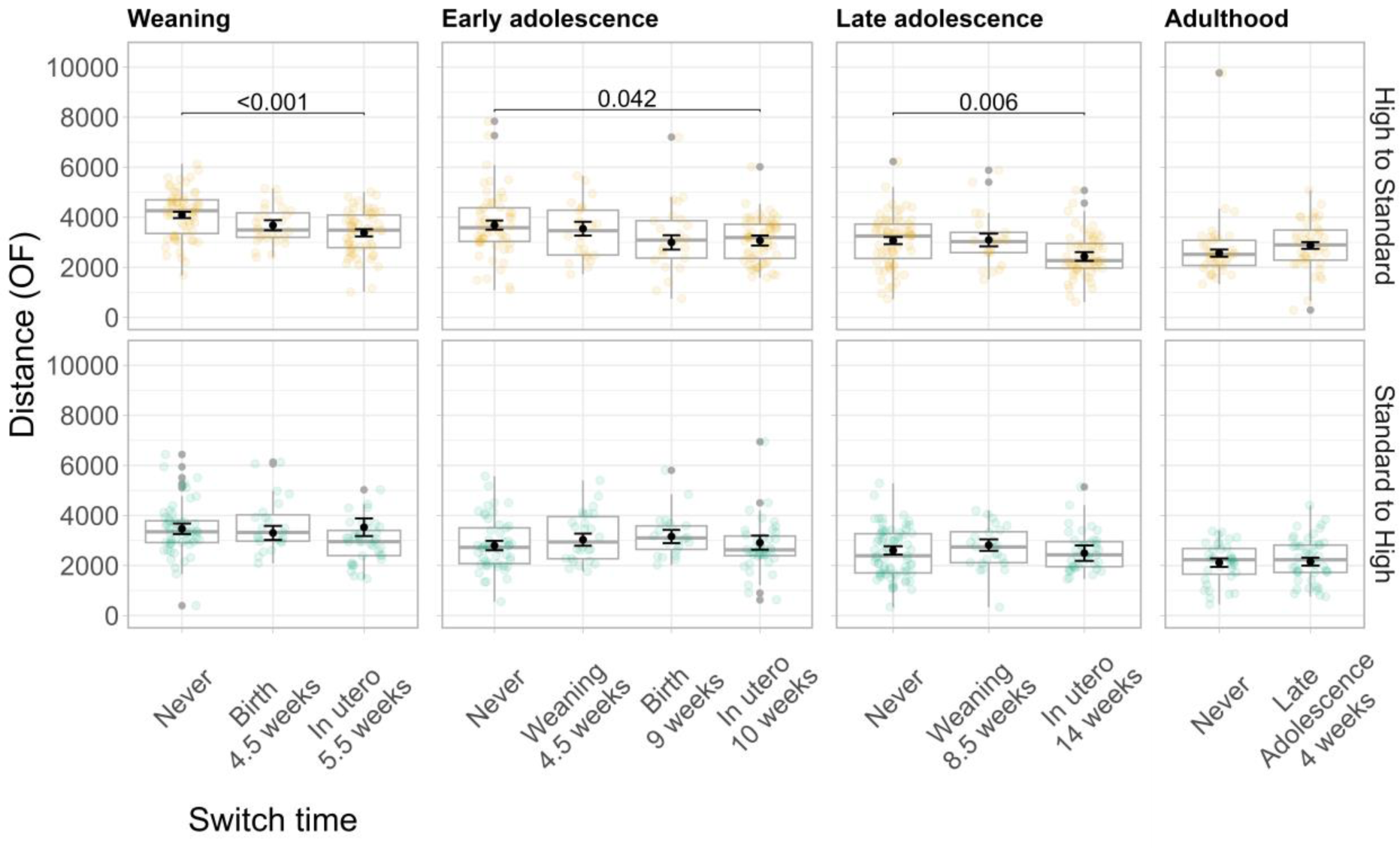
Distance covered in the Open Field across food switch times. The top row shows results comparing mice who have continuously received high-quality food to mice who have been switched from high to standard-quality food. The bottom row shows results comparing mice who have continuously received standard-quality food to mice who have been switched from standard to high-quality food. Each column corresponds to one test age. Within a panel the x-axis denotes different switch times, with never corresponding to the control group. Beneath each food switch time, the number of weeks indicates how long mice have been fed the new food type. The y-axis denotes distance covered in cm. Within a panel boxplots indicate the distribution of the raw data (also shown as lightly colored circles), while point estimates and error bars (in black) indicate model-based estimated marginal means and standard errors. P-values indicate significant differences between mice whose food was switched at different times compared to the control group. P-values have been corrected for multiple post-hoc comparisons with the Tukey procedure.

Our analysis revealed significant effects of switching mice from high-quality to standard-quality food during the mother’s pregnancy, indicating a potential sensitive period in utero (Tables 3-4 and Figure 3 top row). Mice who were switched in utero covered less distance in the Open Field compared to control individuals who continuously received high-quality food. This effect was measurable in weanlings, early adolescents, and late adolescents. Across test ages, we can observe a consistent reduction in distance covered of around 600-700 cm (Table 2). We would like to note that mean differences across all switch- time-groups were not significantly different in early adolescents (Table 4). ANOVA tests whether there are significant mean differences across all groups. However, our contrasts in the regression analysis specifically compare each group mean to the reference group (i.e., the control mice) (Table 3). Our contrasts thus confirm that early adolescent mice who experienced a food switch during the mother’s pregnancy show significantly different behavior compared to control individuals – even though differences across all switch-time- groups measured during early adolescence do not significantly differ.

**Table 2:** Regression output for Distance (OF) across test ages.

|  |  | Kenward-Roger Estimates |  |  |  |  |  | Bootstrapped Estimates |  |  |  |
| --- | --- | --- | --- | --- | --- | --- | --- | --- | --- | --- | --- |
|  | Starting food | *b* | *beta* | *SE* | *df* | *t* | *p* | *b* | 95% CI | *p* | *p adj.* |
| **Weaning** |  |  |  |  |  |  |  |  |  |  |  |
| sex (males) | SQ | -81.98 | -0.04 | 160.87 | 41.80 | -0.51 | 0.613 | -80.68 | (-396.048, 227.811) | 0.597 | 0.838 |
| switch time (in utero) | SQ | 61.69 | 0.03 | 398.67 | 45.43 | 0.15 | 0.878 | 57.46 | (-673.653, 833.272) | 0.875 | 0.875 |
| switch time (birth) | SQ | -164.45 | -0.06 | 307.80 | 60.76 | -0.53 | 0.595 | -162.05 | (-774.872, 427.613) | 0.612 | 0.612 |
| sex (males) | HQ | -56.45 | -0.03 | 149.92 | 141.83 | -0.38 | 0.707 | -54.98 | (-339.972, 232.898) | 0.703 | 0.967 |
| switch time (in utero) | HQ | -714.07 | -0.37 | 191.32 | 20.22 | -3.73 | **0.001** | -706.23 | **(-1088.111, -350.694)** | **<0.001** | **<0.001** |
| switch time (birth) | HQ | -413.23 | -0.17 | 236.45 | 36.26 | -1.75 | 0.089 | -403.33 | (-861.552, 52.786) | 0.081 | 0.162 |
| **Early adolescence** |  |  |  |  |  |  |  |  |  |  |  |

Table 2: Model summary for modeling distance in the open field at different test ages.
|  |  | Kenward-Roger Estimates |  |  |  |  |  | Bootstrapped Estimates |  |  |  |
| --- | --- | --- | --- | --- | --- | --- | --- | --- | --- | --- | --- |
|  | Starting food | *b* | *beta* | *SE* | *df* | *t* | *p* | *b* | 95% CI | *p* | *p adj.* |
| sex (males) | SQ | -137.08 | -0.07 | 180.09 | 60.22 | -0.76 | 0.450 | -136.90 | (-494.485, 224.671) | 0.433 | 0.577 |
| switch time (in utero) | SQ | 115.78 | 0.05 | 338.08 | 25.01 | 0.34 | 0.735 | 112.59 | (-543.494, 740.296) | 0.740 | 0.740 |
| switch time (birth) | SQ | 360.63 | 0.13 | 307.37 | 62.26 | 1.17 | 0.245 | 353.42 | (-232.368, 960.267) | 0.234 | 0.469 |
| switch time (weanling) | SQ | 233.34 | 0.09 | 275.55 | 78.89 | 0.85 | 0.400 | 230.39 | (-314.206, 778.197) | 0.404 | 0.452 |
| sex (males) | HQ | -167.72 | -0.07 | 193.12 | 78.54 | -0.87 | 0.388 | -167.52 | (-540.955, 205.634) | 0.382 | 0.910 |
| switch time (in utero) | HQ | -616.61 | -0.26 | 270.57 | 25.04 | -2.28 | **0.031** | -618.77 | **(-1142.085, -90.985)** | **0.021** | **0.042** |
| switch time (birth) | HQ | -692.19 | -0.21 | 334.05 | 42.76 | -2.07 | **0.044** | -691.48 | **(-1334.853, -43.701)** | **0.037** | 0.147 |
| switch time (weanling) | HQ | -143.79 | -0.04 | 307.00 | 91.30 | -0.47 | 0.641 | -140.16 | (-745.14, 449.417) | 0.656 | 0.848 |
| **Late adolescence** |  |  |  |  |  |  |  |  |  |  |  |
| sex (males) | SQ | 83.29 | 0.05 | 151.24 | 127.12 | 0.55 | 0.583 | 83.61 | (-208.597, 380.67) | 0.572 | 0.763 |
| switch time (in utero) | SQ | -108.04 | -0.05 | 351.52 | 20.47 | -0.31 | 0.762 | -118.01 | (-784.92, 570.899) | 0.727 | 0.727 |
| switch time (weanling) | SQ | 213.90 | 0.09 | 227.05 | 129.64 | 0.94 | 0.348 | 211.75 | (-242.789, 650.011) | 0.359 | 0.741 |
| sex (males) | HQ | -128.44 | -0.06 | 175.16 | 69.42 | -0.73 | 0.466 | -131.28 | (-472.325, 207.836) | 0.468 | 0.623 |
| switch time (in utero) | HQ | -636.53 | -0.31 | 225.50 | 21.43 | -2.82 | **0.010** | -633.98 | **(-1053.94, -189.762)** | **0.003** | **0.006** |
| switch time (weanling) | HQ | 23.56 | 0.01 | 272.41 | 82.83 | 0.09 | 0.931 | 25.13 | (-493.589, 551.409) | 0.932 | 0.932 |
| **Adulthood** |  |  |  |  |  |  |  |  |  |  |  |
| sex (males) | SQ | -115.96 | -0.07 | 183.18 | 77.71 | -0.63 | 0.529 | -116.84 | (-481.724, 238.49) | 0.532 | 0.702 |
| switch time (adult) | SQ | 40.45 | 0.02 | 180.04 | 77.50 | 0.22 | 0.823 | 40.15 | (-306.076, 394.332) | 0.817 | 0.817 |
| sex (males) | HQ | 496.89 | 0.28 | 197.22 | 35.19 | 2.52 | **0.016** | 500.01 | **(112.695, 883.759)** | **0.011** | **0.045** |
| switch time (adult) | HQ | 306.59 | 0.17 | 197.51 | 36.27 | 1.55 | 0.129 | 304.41 | (-79.505, 686.487) | 0.120 | 0.333 |

**Table 3:** Anova summary Distance (OF) across test ages.

Analysis of variance summary with Kenward-Roger degrees of freedom.
| | Starting food | Sum Sq | Mean Sq | NumDF | DenDF | F | p | p adj. | partial $\eta$ | 95% CI |
| --- | --- | --- | --- | --- | --- | --- | --- | --- | --- | --- |
| **Weaning** |  |  |  |  |  |  |  |  |  |  |
| sex | SQ | 126151.49 | 126151 | 1 | 41.80 | 0.26 | 0.613 | 0.836 | 0.007 | (0,1) |
| switch time | SQ | 236271.28 | 118136 | 2 | 55.48 | 0.24 | 0.785 | 0.785 | 0.010 | (0,1) |
| sex | HQ | 106276.37 | 106276 | 1 | 141.83 | 0.14 | 0.707 | 0.944 | 0.001 | (0,1) |
| switch time | HQ | 10649388.84 | 5324694 | 2 | 23.68 | 7.09 | **0.004** | **0.015** | 0.338 | **(0.096,1)** |
| **Early adolescence** |  |  |  |  |  |  |  |  |  |  |
| sex | SQ | 418171.09 | 418171 | 1 | 60.22 | 0.58 | 0.450 | 0.599 | 0.011 | (0,1) |
| switch time | SQ | 1254994.69 | 418332 | 3 | 46.54 | 0.57 | 0.634 | 0.717 | 0.033 | (0,1) |
| sex | HQ | 790926.12 | 790926 | 1 | 78.54 | 0.75 | 0.388 | 0.916 | 0.010 | (0,1) |
| switch time | HQ | 7593299.08 | 2531100 | 3 | 37.39 | 2.39 | 0.084 | 0.168 | 0.212 | (0,1) |
| **Late adolescence** |  |  |  |  |  |  |  |  |  |  |
| sex | SQ | 170316.55 | 170317 | 1 | 127.12 | 0.30 | 0.583 | 0.777 | 0.002 | (0,1) |
| switch time | SQ | 624741.43 | 312371 | 2 | 40.76 | 0.55 | 0.581 | 0.708 | 0.032 | (0,1) |
| sex | HQ | 421805.32 | 421805 | 1 | 69.42 | 0.54 | 0.466 | 0.621 | 0.008 | (0,1) |
| switch time | HQ | 6743048.14 | 3371524 | 2 | 34.85 | 4.25 | **0.022** | **0.045** | 0.231 | **(0.023,1)** |
| **Adulthood** |  |  |  |  |  |  |  |  |  |  |
| sex | SQ | 184082.83 | 184083 | 1 | 77.71 | 0.40 | 0.529 | 0.688 | 0.005 | (0,1) |
| switch time | SQ | 23185.32 | 23185 | 1 | 77.50 | 0.05 | 0.823 | 0.823 | 0.001 | (0,1) |
| sex | HQ | 4448438.11 | 4448438 | 1 | 35.19 | 6.35 | **0.016** | 0.066 | 0.163 | **(0.019,1)** |
| switch time | HQ | 1688661.35 | 1688661 | 1 | 36.27 | 2.41 | 0.129 | 0.334 | 0.067 | (0,1) |

Taken together, these findings suggest that mice are able to adjust their behaviors to a food switch in utero when food quality decreases (from high to standard-quality) but not when it increases. This implies a nutrition-dependent sensitive period in utero. In the appendix we report results from the same analyses for the other three behavioral outcome measures (Tables A6-A17).

### Exploratory analysis: Does the timing of food switch shape changes in behavior across development?

From weaning to early adolescence, the timing of food switch does not shape changes in distance covered in the Open Field when mice were switched from high to standard-quality food (*F*(2,143.89) = .37, *p* = .697; appendix Tables A19-A20). However, switch time significantly shapes changes in distance covered when mice were switched from standard- to high-quality food (*F*(2,110.69) = 4.85, *p* < .001; appendix Tables A19-20). Post-hoc comparisons reveal that only control individuals who consistently received standard-quality food significantly start to cover less distance as they become older (estimated difference = 721.17, *t*(112.82) = 6.03, *p* < .001; appendix Tables A21-A22). In contrast, mice who experienced a food switch in utero (estimated difference = 203.16, *t*(108.60) = 1.52, *p* = .132) or at birth (estimated difference = 257.46, *t*(110.86) = 1.44, *p* = .153) do not significantly differ from each other in distance covered. We illustrate this relationship in the appendix in Figure A4. Our results may indicate that switching mice to a more nutritious diet slows the age-related transition from active to passive stress coping.

From early to late adolescence, switch time does not shape changes in distance covered in the Open Field, irrespective of diet (high-quality food: *F*(2,143.31) = .37, *p* = .692 and standard-quality food: *F*(2,119.74) = .77, *p* = .622; appendix Tables A23-A24). In the appendix we report results from the same analyses for the other three behavioral outcome measures (Tables A19-A24).

### A sensitive period for stress-coping during the fetal stage?

Figure 3 shows the distribution of mice who have experienced a food switch in utero (in either direction) across the measured test ages. For comparison, we added the distributions of the respective control groups. This simple descriptive figure illustrates three key patterns that emerged from our analyses. First, it highlights that distance covered in the Open Field tends to decrease with age. Second, it shows how mice who have been switched from high to standard-quality food appear to adopt different behaviors in the Open Field compared to control individuals. In contrast, mice who have been switched from standard to high-quality food appear to remain similar to the control group. Third, the figure illustrates how a switch from standard to high-quality food might slow the age-related decline of distance covered in the Open Field.

**Figure 3:**
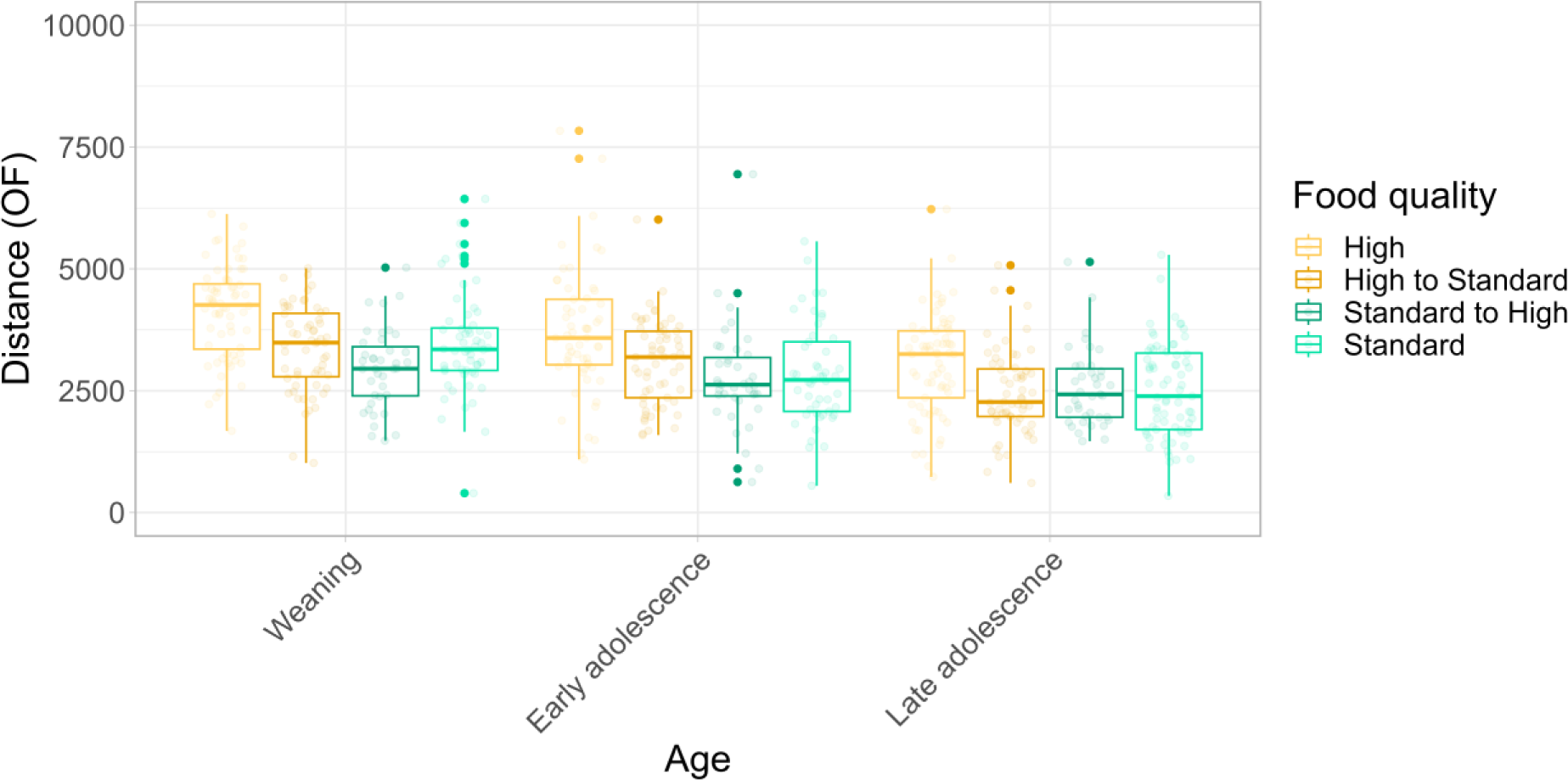
Distance covered in the Open Field across development in mice who experienced a food switch in utero. For comparison purposes, the figure additionally shows data from control individuals. The x-axis denotes test ages and the y-axis distance covered in cm. For each test age, boxplots indicate the distribution of the raw data for both control (mice who continuously received one food type) and treatment individuals (mice who experienced a food switch in utero). Yellow corresponds to mice who started with standard-quality food and green to mice who started with high-quality food. Lighter shades of yellow and green correspond to control individuals and darker shades to treatment individuals.

## Discussion

### The ontogeny of stress coping

We found that stress coping changes across ontogeny in (cage-housed) mice, even when there is no experimental manipulation. As mice get older, they develop increasingly more passive stress-coping styles, indicating higher levels of risk-aversion (Krebs et al., 2019). At the age of 17 weeks, animals in our study expressed comparable trait values to studies investigating stress-coping behavior in adult wild house mice living under cage or semi-natural housing conditions (Küçüktaş & Guenther, 2022; Prabh et al., 2023). Understanding the ontogeny of stress-coping lays the foundation for studying how and when experiences can alter developmental trajectories, and when we can measure their effects.

### A nutrition-dependent sensitive period for stress-coping

We observed that mice experiencing decreases in nutritional conditions in utero adopt passive stress-coping strategies, implying higher levels of risk aversion. These effects are measurable in weanlings and across adolescence. Mice might employ these strategies as an insurance against variation in food quality (Mathot et al., 2012). Mice previously receiving high-quality food have reproductive value to lose and might protect these ‘assets’ by reducing risky exposure to threats and predators (Clark, 1994; Lopez-Hervas et al., 2024; McElreath et al., 2007; Wolf et al., 2007).

Our findings dovetail with prior work documenting nutrition-dependency in the capacity of mice to adjust stress-coping and life-history traits across one generation (Lopez-Hervas et al., 2024). Across both studies, effects are stronger (or only present) in mice who were switched away from calorie-dense, high-quality food. This implies that calorie-dense food might be necessary to supply energy for plasticity. However, in contrast to the present study, mice whose parents’ were switched from high- to standard-quality food showed more active stress-coping and thus higher risk-taking tendencies compared to controls (Lopez-Hervas et al., 2024). Below we will discuss the differences between these studies.

Additionally, our data suggest nutrition-dependent effects on age-related changes in stress coping. In our sample, experiencing increases in nutritional conditions seemed to slow the age-related transition towards passive stress coping. Mice who were switched to high-quality food reduced active stress coping to a lesser extent compared to controls. However, we note that both treatment and control mice have similar trait values in late adolescence. The difference in trajectories appears to be driven by the different starting points: Weanlings who were switched to a high-quality diet show descriptively lower levels of active stress coping compared to controls. This difference was not statistically significant. It is unclear whether this descriptive difference reflects a true difference resulting from the treatment or simply noise. To reliably estimate whether the timing of food switch alters developmental trajectories of stress coping future studies with larger sample sizes are needed.

Behavioral adjustments to changes in food quality are likely mediated through (neuro-) physiological changes. Previous work in wild house mice has shown that a low-protein diet in utero prompts rapid post-natal growth and reduced longevity in male mice, without affecting neural development (Buchanan et al., 2022). It is difficult to directly relate these findings to our study, as we have not contrasted high- and low-quality diets or measured neural development. Future studies, may consider using standard-quality food as a control diet with high- and low-quality food as treatment conditions. Additionally, such studies would ideally monitor neural and hormonal development to provide further insights into the mechanistic pathways underlying sensitive periods for stress coping (Trillmich et al., 2018).

### A role for fetal programming

Our data suggest a potential role for fetal programming of stress-coping behaviors. Empirical evidence in human and non-human animals suggests that nutritional conditions in utero can shape later-life health and well-being (Horton, 2005; Jones, 2005; Wells & Johnstone, 2017). The underlying idea is that the mother provides cues, integrating over a lifetime of her experiences, which the developing fetus uses to ‘predict’ the long-term expected nutritional environment (Kuzawa, 2005; Kuzawa & Thayer, 2011). However, fetal programming assumes that information provided through the mother better predicts expected nutritional conditions than information sampled postnatally (so called external predictive adaptive response; Nettle et al., 2013). This assumption holds more likely when environmental conditions are stable (or autocorrelated) across generations. When organisms are adapted to expect changes across generations, they should rely on personal information instead (Guenther et al., 2021). An external predictive adaptive response explanation seems plausible in house mice living in human-made environments. These environments dampen seasonal variation in temperature, food-availability and precipitation, implying higher stability and predictability.

However, two limitations of our experimental design warrant caution in drawing certain conclusions about a sensitive period in utero. First, we do not have a full-factorial design. For example, we did not measure late adolescent mice who experienced a food switch at birth and we also do not have measures of adult mice who experienced food switches in utero or at birth. Second, the number of mice who experienced a food switch at birth is much smaller compared to those experiencing a food switch in utero (see Figure 1). The reason for this is that mothers of mice who were switched from high to standard-quality food at birth withdrew maternal investment and ate their offspring. Thus, we may not have enough power to detect significant effects of food switches during pregnancy. At least in early adolescence the descriptive differences between control and treatment mice are similar for the in utero and at birth condition. Thus, while our data clearly point towards an early sensitive period for the effects of changes in food quality, we cannot (yet) clearly delineate its timing and duration. Future studies are needed to clearly disentangle whether effects are stronger for food switches in utero or at birth.

### Should we always expect sensitive periods early in life?

Across species and traits interest has grown in sensitive periods at later developmental stages, such as adolescence. This pattern appears to be prominent for the development of social traits in mammals (reviewed in Walasek et al., n.d.). Modeling work has suggested that we might expect sensitive periods later in ontogeny, when the value of information from cues or the environment itself changes across ontogeny (Walasek et al., 2022b, 2022a).

Manipulating changes in nutritional quality at several time points during ontogeny, would allow us to test whether the sensitive period for stress coping could shift depending on the rate of environmental change.

### Difference between mice living in semi-natural conditions and cage-housed mice

Our study is closely related to two recent studies exploring how nutritional conditions shape stress-coping and life-history traits in wild house-mice living in semi-natural enclosures (Lopez-Hervas et al., 2024; Prabh et al., 2023). Both studies report that mice who received high-quality food develop passive, risk-averse stress-coping strategies. The authors argue that these patterns are in line with asset-protection theory: Mice who stand more to lose, should take fewer risks and protect their assets (Clark, 1994; Wolf et al., 2007). These findings resonate with a recent study in African clawed frog tadpoles (*Xenopus laevis*) (Beyts et al., 2023). Tadpoles who experienced food restriction early in life took greater foraging risks in novel environments compared to food-unrestricted tadpoles. In cage-housed mice, we observe the opposite pattern, with mice continuously receiving high-quality food developing more active, risk-seeking coping strategies. From a theoretical point of view, we could argue that high-quality food increases the chances of mice successfully taking risks (Barclay et al., 2018). A study in hihis (*Notiomystis cincta*) lends support to this idea (Richardson et al., 2019). It documented weak, positive associations between early-life carotenoid supplementation and boldness, which tends to be positively associated with risk taking (Eccard et al., 2022).

Even if we can find theoretical explanations for each study in isolation, how can we reconcile these contrasting patterns within the same species and even the same population of mice (our mice being direct descendants of those from the two previously mentioned studies)? One crucial difference between mice living in cages and those in semi-natural enclosures is the social environment. Cage-housed mice only live in same-sex pairs, whereas enclosure-housed mice live in groups of 80-140. Thus, one stark difference between the study designs is that cage-housed mice are not able to reproduce, potentially providing them with more energy to invest into exploration.

However, the social environment is not only important for providing reproductive opportunities. Previous work has highlighted the importance of the social environment for the development of behavior and personality (Eccard & Herde, 2013; MacLeod et al., 2023; Sachser et al., 2013). For example, a series of studies in guinea pigs suggest that males who have been housed in groups during adolescence develop to be less aggressive adults compared to males housed in isolation (reviewed in Sachser et al., 2020). Living in social groups might also help individuals to better recover from stressful experiences (Kikusui et al., 2006; Komdeur & Ma, 2021). In contrast to these beneficial effects of social groups, experiencing higher population density could also imply larger competition for resources, potentially making individuals more protective of their assets. Future work could explicitly study the role of the social environment by varying group size in captive and (semi-)free living mice. In the best-case scenario the stark differences across studies point to the social environment as a potential moderator for the development of stress coping. In the worst-case scenario, these results call the validity of studies in captive mice and potentially other animals into question (Kondrakiewicz et al., 2019) as has been discussed in the literature for a while (Fisher et al., 2015; Niemelä & Dingemanse, 2014). More work is needed to understand the effects of laboratory housing conditions on experimental findings.

Throughout this discussion section we have suggested several ways in which future work could clarify some of the questions raised by our study. These studies would shed light on the mechanisms underlying stress-coping behaviors in rodents, providing important insights into how and when organisms are able to adapt to changing environmental conditions.

## Acknowledgements

We thank Fragkiskos Darmis and Willem Frankenhuis for valuable feedback on earlier versions of this manuscript.

## Author Contributions

AG conceived of the study and MJ collected the data. NW analyzed the data and wrote the manuscript. AG and NW revised the manuscript.

## Data Availability Statement

We will upload the data to the OSF project page ([project page](https://osf.io/v7drq/?view_only=b4eda60ec8394e38ba27940459b97b4c)) upon publication. This is currently a view-only link to maintain anonymity of the authors for peer review.

## Ethics

Housing of wild mice is approved and regularly controlled by the Veterinäramt Plön under licence: 1401-144/PLÖ-004697. All animals were handled, and procedures were carried out according to national guidelines. For animals tested in the Open Field and Elevated Plus Maze tests, experiments were performed under licences V244 – 31223/2019(62-5/19).

