## Appendix for "Personality development in wild house mice: Evidence for a nutrition-dependent sensitive period early in life"

Legend

| | |
|---|---|
| Distance (OF) / distance_of | Distance covered in the Open Field (in cm) |
| Center (OF) / center_of | Time spent in the center of the open field (in percent) |
| Distance (EP) / distance_ep | Distance covered in the Elevated Plus Maze (in cm) |
| Brightarm (EP) / brightarm_ep | Time spent in the bright arms of the Elevated Plus Maze (in percent) |
| Sex | Sex of the mouse |
| startingfood | The food on which mice started the experiment (both in control and treatment individuals); can either be SQ (= standard quality) or HQ (= high quality) |
| testage | The age at which mice were tested in the Open Field and the Elevated Plus Maze. The levels correspond to 5week (~ weaning), 9week (~ early adolescence), 13week (~late adolescence), and 17week (~ adulthood), indicating the respective age in weeks. |
| Switch_time | The time during ontogeny at which treatment mice experienced a food switch. The levels include (the mother's) pregnancy, birth, weanling, and adult (corresponding to a food switch at late adolescence). |

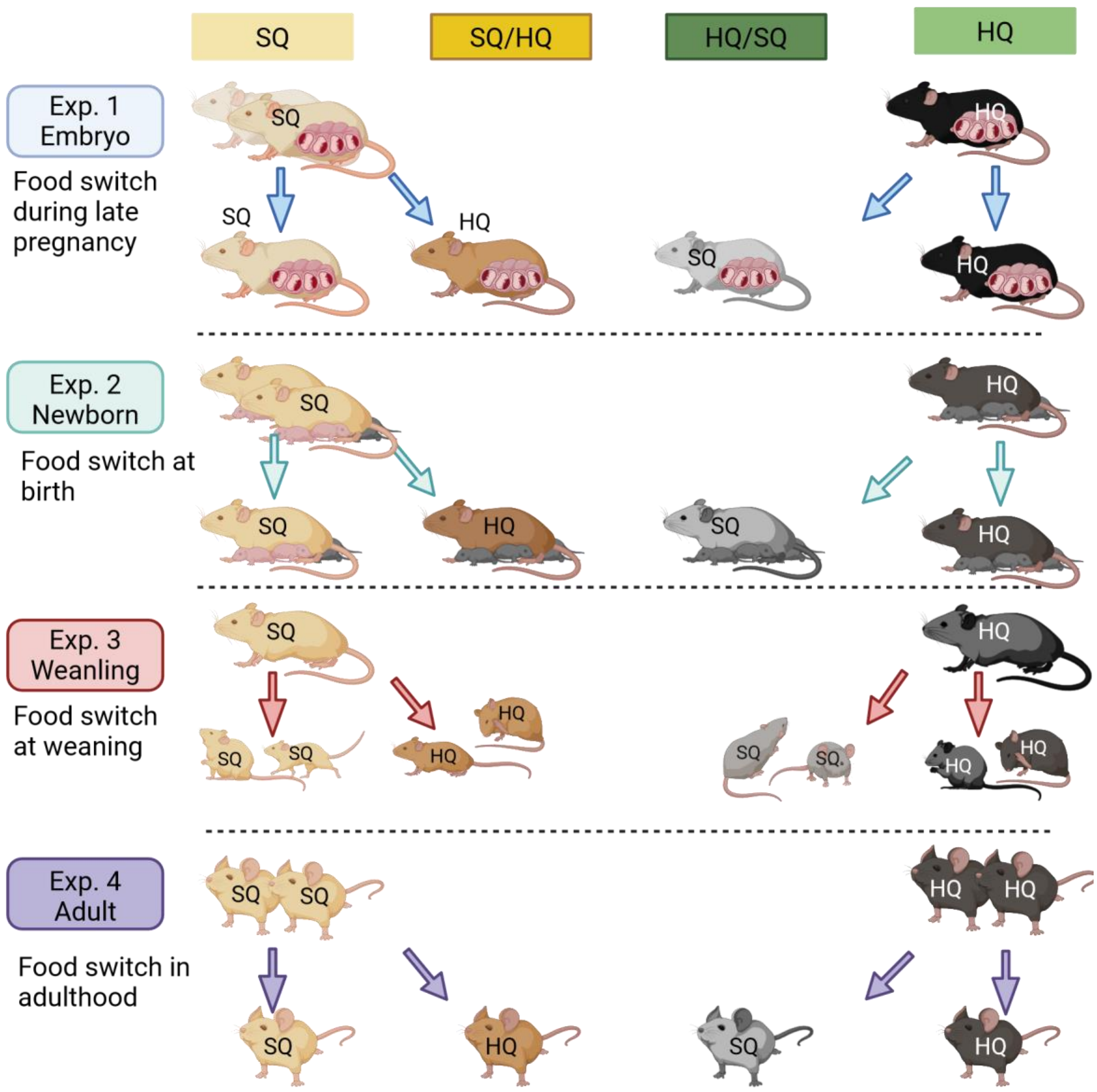


*Figure A1:* Overview over the experimental design. This figure illustrates four experiments, each of which introduces a food switch during a different ontogenetic time point. Within each experiment there are two control groups of mice who continuously receive standard quality (SQ) or high quality (HQ) food. Additionally, each experiment contains two treatment groups that experience a food switch either from standard to high quality food (SQ/HQ) or the other way round (HQ/SQ).

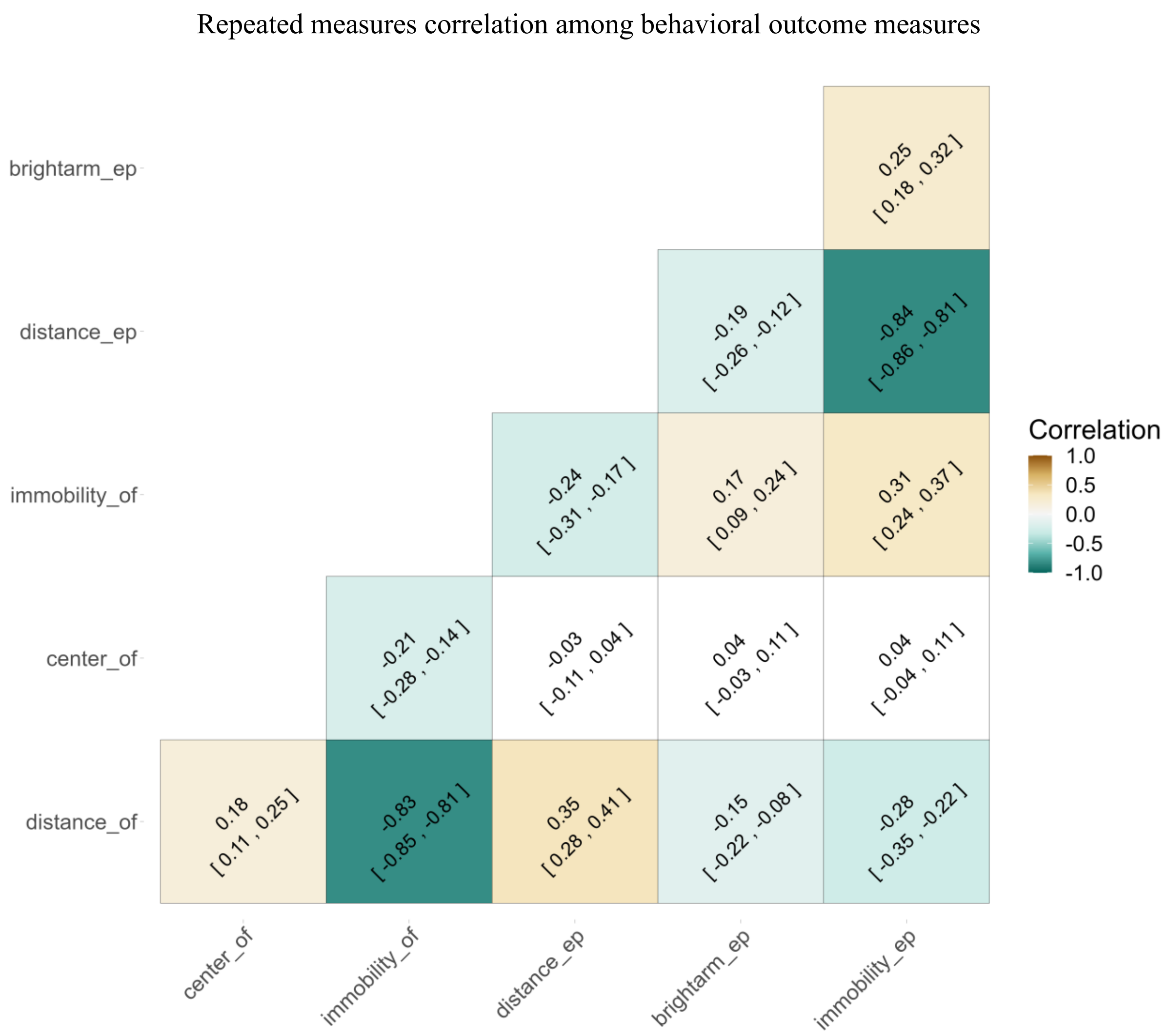


*Figure A2:* Repeated measures correlations among behavioral outcome measures. Each cell corresponds to one repeated measures correlation between the measures indicates by the row and column. Colored cells indicate significant correlations at an alpha-level of .005. Color intensity is proportional to the strength of the correlation coefficient.

#### Results analyses in control individuals (SQ, HQ without food switch)

**Table A1. Regression output for all outcome variables.**

| | Kenward-Roger Estimates | | | | | Bootstrapped Estimates | | | |
|---|---|---|---|---|---|---|---|---|---|
| | **Estimate** | ***SE*** | **df** | ***t*** | ***p*** | **Estimate** | **95% CI** | ***p*** | ***p adj.*** |
| **distance (OF)** | | | | | | | | | |
| sex2 | -33.22 | 157.35 | 68.61 | -0.21 | 0.833 | -30.39 | (-339.494, 273.355) | 0.842 | 0.842 |
| starting_food1 | 407.20 | 98.04 | 73.66 | 4.15 | **<0.001** | 408.22 | (213.332, 595.999) | **<0.001** | **<0.001** |
| testage1 | -294.50 | 79.39 | 308.44 | -3.71 | **<0.001** | -291.82 | (-448.094, -134.74) | **<0.001** | **<0.001** |
| testage2 | -536.03 | 103.84 | 348.44 | -5.16 | **<0.001** | -536.58 | (-745.128, -337.046) | **<0.001** | **<0.001** |
| testage3 | 672.91 | 87.62 | 346.48 | 7.68 | **<0.001** | 671.98 | (502.151, 845.613) | **<0.001** | **<0.001** |
| starting_food1:testage1 | -53.01 | 79.36 | 309.63 | -0.67 | 0.505 | -51.58 | (-210.839, 103.151) | 0.510 | 0.826 |
| starting_food1:testage2 | 225.24 | 104.12 | 351.26 | 2.16 | **0.031** | 227.22 | (23.67, 434.393) | **0.028** | 0.110 |
| starting_food1:testage3 | -164.26 | 87.69 | 349.00 | -1.87 | 0.062 | -166.59 | (-333.678, 9.34) | 0.063 | 0.251 |
| **center (OF)** | | | | | | | | | |
| sex2 | 0.17 | 0.12 | 77.46 | 1.44 | 0.153 | 0.18 | (-0.05, 0.41) | 0.135 | 0.270 |
| starting_food1 | 0.13 | 0.06 | 53.42 | 1.97 | 0.054 | 0.12 | (0, 0.25) | **0.041** | 0.081 |
| testage1 | -0.16 | 0.07 | 313.80 | -2.34 | **0.020** | -0.16 | (-0.29, -0.03) | **0.022** | **0.043** |
| testage2 | 0.30 | 0.09 | 352.45 | 3.52 | **<0.001** | 0.30 | (0.13, 0.47) | **<0.001** | **<0.001** |
| testage3 | 0.04 | 0.07 | 349.95 | 0.59 | 0.557 | 0.04 | (-0.1, 0.19) | 0.566 | 0.566 |
| starting_food1:testage1 | 0.01 | 0.07 | 315.26 | 0.19 | 0.852 | 0.01 | (-0.12, 0.14) | 0.826 | 0.826 |
| starting_food1:testage2 | 0.13 | 0.09 | 354.65 | 1.45 | 0.147 | 0.13 | (-0.04, 0.3) | 0.138 | 0.185 |
| starting_food1:testage3 | -0.06 | 0.07 | 351.56 | -0.85 | 0.397 | -0.06 | (-0.2, 0.08) | 0.382 | 0.510 |
| **distance (EP)** | | | | | | | | | |
| sex2 | -102.18 | 214.95 | 83.09 | -0.48 | 0.636 | -97.76 | (-499.194, 312.989) | 0.644 | 0.842 |
| starting_food1 | 197.77 | 110.73 | 54.35 | 1.79 | 0.080 | 198.68 | (-7.702, 416.28) | 0.060 | 0.081 |
| testage1 | -93.92 | 88.79 | 295.14 | -1.06 | 0.291 | -96.49 | (-268.555, 77.271) | 0.271 | 0.328 |
| testage2 | -363.49 | 116.67 | 326.65 | -3.12 | **0.002** | -364.60 | (-588.498, -132.567) | **<0.001** | **<0.001** |
| testage3 | 226.90 | 99.17 | 328.88 | 2.29 | **0.023** | 225.12 | (30.389, 424.961) | **0.021** | **0.028** |
| starting_food1:testage1 | 40.80 | 88.86 | 296.51 | 0.46 | 0.646 | 40.92 | (-127.304, 220.007) | 0.640 | 0.826 |
| starting_food1:testage2 | 179.81 | 117.05 | 329.11 | 1.54 | 0.125 | 180.48 | (-40.925, 403.693) | 0.118 | 0.185 |
| starting_food1:testage3 | -111.12 | 99.29 | 331.25 | -1.12 | 0.264 | -111.25 | (-302.266, 80.904) | 0.250 | 0.499 |
| **brightarm (EP)** | | | | | | | | | |
| sex2 | -3.93 | 1.64 | 69.12 | -2.39 | **0.019** | -3.93 | (-7.091, -0.632) | **0.019** | 0.077 |
| starting_food1 | -0.28 | 0.84 | 77.91 | -0.33 | 0.740 | -0.27 | (-1.897, 1.377) | 0.750 | 0.750 |
| testage1 | 1.11 | 1.11 | 318.82 | 1.00 | 0.316 | 1.09 | (-1.091, 3.304) | 0.328 | 0.328 |
| testage2 | 3.36 | 1.39 | 350.42 | 2.41 | **0.017** | 3.36 | (0.574, 6.193) | **0.017** | **0.017** |
| testage3 | -4.40 | 1.19 | 350.62 | -3.69 | **<0.001** | -4.37 | (-6.699, -2.095) | **<0.001** | **<0.001** |
| starting_food1:testage1 | -1.99 | 1.11 | 319.85 | -1.79 | 0.074 | -1.94 | (-4.166, 0.195) | 0.072 | 0.288 |
| starting_food1:testage2 | 0.70 | 1.40 | 353.39 | 0.50 | 0.617 | 0.70 | (-1.908, 3.451) | 0.614 | 0.614 |

**Table A1. Regression output for all outcome variables.**

| | Kenward-Roger Estimates | | | | | Bootstrapped Estimates | | | |
|---|---|---|---|---|---|---|---|---|---|
| | **Estimate** | ***SE*** | **df** | ***t*** | ***p*** | **Estimate** | **95% CI** | ***p*** | ***p adj.*** |
| starting_food1:testage3 | 0.09 | 1.19 | 351.96 | 0.07 | 0.942 | 0.07 | (-2.257, 2.358) | 0.955 | 0.955 |

*Table A1: Model summary for all outcome variables. All variables were fitted with mixed effects models. We attempted to fit each model with nested random intercepts with mice nested in cages and cages in families. If the model did not converge, we simplified the random effects structure (see Table A5). We applied a square root transformation to Center (OF) because the distribution of times spent in the center was skewed towards the right. For all mixed models degrees of freedom were computed with the Kenward-Roger approximation. Additionally, we computed bootstrapped parameters estimates, confidence intervals, and p-values. Per model, we applied a correction for multiple testing (FDR correction) to these bootstrapped p-values across the four outcome variables which use the same set of predictors. Starting food and test age were sum-to-zero coded to avoid multicollinearity resulting from their involvement in an interaction. Sex was dummy coded with females as the reference group. Center (OF) and brightarm (EP) are measured in percent.*

**Table A2. Anova summary**

| | **Sum Sq** | **Mean Sq** | **NumDF** | **DenDF** | ***F*** | ***p*** | ***p adj.*** |
|---|---|---|---|---|---|---|---|
| **distance (OF)** | | | | | | | |
| sex | 42354.99 | 42354.99 | 1 | 68.61 | 0.04 | 0.833 | 0.833 |
| starting_food | 16395267.95 | 16395267.95 | 1 | 73.66 | 17.25 | **<0.001** | **<0.001** |
| testage | 64724458.45 | 21574819.48 | 3 | 321.25 | 22.70 | **<0.001** | **<0.001** |
| starting_food:testage | 5815645.08 | 1938548.36 | 3 | 322.73 | 2.04 | 0.108 | 0.433 |
| **center (OF)** | | | | | | | |
| sex | 1.44 | 1.44 | 1 | 77.46 | 2.08 | 0.153 | 0.306 |
| starting_food | 2.67 | 2.67 | 1 | 53.42 | 3.87 | 0.054 | 0.106 |
| testage | 12.80 | 4.27 | 3 | 325.11 | 6.18 | **<0.001** | **<0.001** |
| starting_food:testage | 1.71 | 0.57 | 3 | 326.27 | 0.83 | 0.480 | 0.480 |
| **distance (EP)** | | | | | | | |
| sex | 264699.08 | 264699.08 | 1 | 83.09 | 0.23 | 0.636 | 0.833 |
| starting_food | 3736988.34 | 3736988.34 | 1 | 54.35 | 3.19 | 0.080 | 0.106 |
| testage | 15005638.50 | 5001879.50 | 3 | 304.74 | 4.27 | **0.006** | **0.006** |
| starting_food:testage | 3507878.34 | 1169292.78 | 3 | 306.12 | 1.00 | 0.394 | 0.480 |
| **brightarm (EP)** | | | | | | | |
| sex | 1128.36 | 1128.36 | 1 | 69.12 | 5.73 | **0.019** | 0.078 |
| starting_food | 21.81 | 21.81 | 1 | 77.91 | 0.11 | 0.740 | 0.740 |
| testage | 2931.24 | 977.08 | 3 | 326.18 | 4.96 | **0.002** | **0.003** |
| starting_food:testage | 690.24 | 230.08 | 3 | 327.23 | 1.17 | 0.322 | 0.480 |

*Table A2: Analysis of variance summary with Kenward-Roger degrees of freedom. Per model, we applied a correction for multiple testing (FDR correction) across the four outcome variables which use the same set of predictors. Center (OF) and brightarm (EP) are measured in percent.*

**Table A3. Posthoc pairwise comparisons.**

| | Tukey procedure | | | | | | Unadjusted bootstrapped | |
|---|---|---|---|---|---|---|---|---|
| | **Estimate** | ***SE*** | **df** | ***t*** | **95% CI** | ***p*** | **Estimate** | **95% CI** |
| **distance (OF)** | | | | | | | | |
| HQ - SQ | 814.40 | 196.08 | 73.66 | 4.15 | **(423.666, 1205.125)** | **<0.001** | 816.44 | (426.663, 1191.997) |
| 5week - 9week | 515.29 | 133.97 | 295.49 | 3.85 | **(169.159, 861.419)** | **<0.001** | 514.53 | (247.125, 783.847) |
| 13week - 5week | -967.42 | 135.12 | 345.87 | -7.16 | **(-1316.245, -618.591)** | **<0.001** | -963.89 | (-1231.817, -696.85) |
| 13week - 9week | -452.13 | 136.13 | 330.94 | -3.32 | **(-803.639, -100.618)** | **0.005** | -450.42 | (-720.694, -179.232) |
| 13week - 17week | 241.53 | 145.05 | 297.99 | 1.67 | (-133.226, 616.288) | 0.344 | 243.96 | (-51.333, 537.519) |
| 17week - 5week | -1208.95 | 164.70 | 360.61 | -7.34 | **(-1634.038, -783.859)** | **<0.001** | -1206.74 | (-1539.642, -894.826) |
| 17week - 9week | -693.66 | 166.10 | 351.98 | -4.18 | **(-1122.408, -264.911)** | **<0.001** | -693.72 | (-1016.038, -375.364) |
| **center (OF)** | | | | | | | | |

**Table A3. Posthoc pairwise comparisons.**

| | Tukey procedure | | | | | | Unadjusted bootstrapped | |
|---|---|---|---|---|---|---|---|---|
| | Estimate | *SE* | df | *t* | 95% CI | *p* | Estimate | 95% CI |
| 5week - 9week | 0.90 | 0.44 | 124.19 | 2.06 | (-0.237, 2.047) | 0.171 | 0.90 | (0.057, 1.773) |
| 13week - 5week | -0.76 | 0.44 | 111.12 | -1.73 | (-1.913, 0.387) | 0.313 | -0.75 | (-1.651, 0.104) |
| 13week - 9week | 0.14 | 0.42 | 111.12 | 0.34 | (-0.949, 1.232) | 0.987 | 0.15 | (-0.672, 0.956) |
| 13week - 17week | -1.96 | 0.54 | 111.12 | -3.65 | **(-3.366, -0.562)** | **0.002** | -1.96 | (-3.053, -0.922) |
| 17week - 5week | 1.20 | 0.60 | 124.19 | 1.99 | (-0.369, 2.77) | 0.197 | 1.21 | (0.041, 2.421) |
| 17week - 9week | 2.11 | 0.59 | 137.19 | 3.59 | **(0.579, 3.632)** | **0.003** | 2.10 | (0.98, 3.261) |
| **distance (EP)** | | | | | | | | |
| 5week - 9week | -3.62 | 151.60 | 283.60 | -0.02 | (-395.404, 388.162) | 1.000 | -5.80 | (-301.116, 295.236) |
| 13week - 5week | -320.82 | 152.57 | 327.93 | -2.10 | (-714.792, 73.161) | 0.154 | -322.05 | (-618.699, -18.768) |
| 13week - 9week | -324.44 | 154.54 | 311.13 | -2.10 | (-723.598, 74.724) | 0.156 | -328.09 | (-628.471, -17.508) |
| 13week - 17week | 269.57 | 160.95 | 283.16 | 1.67 | (-146.382, 685.525) | 0.339 | 269.47 | (-46.707, 573.251) |
| 17week - 5week | -590.39 | 186.04 | 340.21 | -3.17 | **(-1070.691, -110.083)** | **0.009** | -590.99 | (-951.781, -227.195) |
| 17week - 9week | -594.01 | 188.31 | 329.18 | -3.15 | **(-1080.273, -107.743)** | **0.009** | -593.06 | (-967.287, -211.856) |
| **brightarm (EP)** | | | | | | | | |
| 5week - 9week | -4.33 | 1.93 | 299.53 | -2.24 | (-9.309, 0.658) | 0.114 | -4.28 | (-8.057, -0.618) |
| 13week - 5week | 5.51 | 1.82 | 351.53 | 3.02 | **(0.803, 10.216)** | **0.014** | 5.50 | (1.822, 9.096) |
| 13week - 9week | 1.18 | 1.88 | 338.52 | 0.63 | (-3.68, 6.048) | 0.923 | 1.18 | (-2.541, 4.82) |
| 13week - 17week | -2.25 | 2.04 | 308.69 | -1.10 | (-7.512, 3.016) | 0.688 | -2.27 | (-6.401, 1.872) |
| 17week - 5week | 7.76 | 2.18 | 354.42 | 3.56 | **(2.13, 13.386)** | **0.002** | 7.74 | (3.539, 12.184) |
| 17week - 9week | 3.43 | 2.24 | 350.62 | 1.53 | (-2.34, 9.204) | 0.418 | 3.47 | (-0.906, 7.675) |

*Table A3: Posthoc pairwise comparisons of significant categorical predictors. Confidence intervals and p-values have been adjusted for multiple comparisons using the Tukey procedure. The last two columns report bootstrapped estimates and confidence intervals. It is important to note that these bootstrapped confidence intervals are not adjusted for multiple testing. Center (OF) and brightarm (EP) are measured in percent.*

**Table A4. Estimated marginal means and medians.**

| | Marginal means | | | | Bootstrapped marginal medians | |
|---|---|---|---|---|---|---|
| | Mean | *SE* | df | 95% CI | Median | 95% CI |
| **distance (OF)** | | | | | | |
| SQ | 2705.06 | 150.53 | 52.89 | (2403.131, 3006.997) | 2701.82 | (2405.946, 2992.55) |
| HQ | 3519.46 | 142.66 | 50.57 | (3232.995, 3805.925) | 3517.60 | (3243.179, 3796.773) |
| 5week | 3785.18 | 136.54 | 88.23 | (3513.85, 4056.502) | 3782.80 | (3512.874, 4044.975) |
| 9week | 3269.89 | 141.55 | 97.58 | (2988.976, 3550.798) | 3265.08 | (2987.534, 3542.007) |
| 13week | 2817.76 | 129.97 | 77.73 | (2559, 3076.517) | 2815.63 | (2564.074, 3076.038) |
| 17week | 2576.23 | 156.93 | 138.19 | (2265.933, 2886.522) | 2572.99 | (2260.815, 2877.91) |
| **center (OF)** | | | | | | |
| SQ | 3.58 | 0.36 | 111.12 | (2.875, 4.289) | 3.60 | (2.938, 4.316) |
| HQ | 4.64 | 0.39 | 124.19 | (3.878, 5.404) | 4.65 | (3.946, 5.424) |
| 5week | 4.23 | 0.39 | 124.19 | (3.461, 4.996) | 4.23 | (3.505, 5.019) |
| 9week | 3.32 | 0.36 | 137.19 | (2.609, 4.039) | 3.33 | (2.663, 4.094) |

**Table A4. Estimated marginal means and medians.**

| | Marginal means | | | | Bootstrapped marginal medians | |
|---|---|---|---|---|---|---|
| | **Mean** | ***SE*** | **df** | **95% CI** | **Median** | **95% CI** |
| 13week | 3.47 | 0.33 | 111.12 | (2.812, 4.119) | 3.48 | (2.857, 4.16) |
| 17week | 5.43 | 0.53 | 195.02 | (4.381, 6.477) | 5.45 | (4.462, 6.532) |
| **distance (EP)** | | | | | | |
| SQ | 3312.64 | 165.15 | 42.41 | (2979.439, 3645.838) | 3310.62 | (2995.781, 3640.647) |
| HQ | 3708.19 | 153.37 | 38.50 | (3397.826, 4018.544) | 3712.73 | (3417.964, 4002.431) |
| 5week | 3737.31 | 147.00 | 72.27 | (3444.281, 4030.334) | 3739.53 | (3456.419, 4017.304) |
| 9week | 3740.93 | 153.55 | 82.56 | (3435.495, 4046.362) | 3743.61 | (3452.077, 4051.802) |
| 13week | 3416.49 | 140.01 | 74.72 | (3137.553, 3695.43) | 3416.99 | (3145.41, 3683.867) |
| 17week | 3146.92 | 171.16 | 139.36 | (2808.518, 3485.322) | 3147.93 | (2810.581, 3474.798) |
| **brightarm (EP)** | | | | | | |
| SQ | 48.97 | 1.22 | 73.42 | (46.545, 51.402) | 48.97 | (46.517, 51.329) |
| HQ | 48.42 | 1.14 | 79.61 | (46.152, 50.681) | 48.41 | (46.25, 50.628) |
| 5week | 44.30 | 1.40 | 298.98 | (41.545, 47.048) | 44.33 | (41.554, 47.079) |
| 9week | 48.62 | 1.50 | 305.47 | (45.68, 51.564) | 48.62 | (45.668, 51.505) |
| 13week | 49.81 | 1.30 | 226.70 | (47.251, 52.361) | 49.79 | (47.243, 52.341) |
| 17week | 52.05 | 1.74 | 293.69 | (48.636, 55.473) | 52.08 | (48.527, 55.518) |

*Table A4: Estimated marginal means for all categorical predictors. The last two columns indicate bootstrapped marginal medians and confidence intervals. Center (OF) and brightarm (EP) are measured in percent.*

**Table A5. Meta information for models in control individuals**

| | n | fullid:(cage:family) | cage:family | fullid:cage | family | cage | ICC_adj | ICC_unadj | r2_m | r2_c |
|---|---|---|---|---|---|---|---|---|---|---|
| **distance (OF)** | 440 | 172 | 96 | | 45 | | 0.354 | 0.234 | 0.201 | 0.484 |
| **center (OF)** | 440 | 172 | 96 | | 45 | | 0.236 | 0.124 | 0.060 | 0.282 |
| **distance (EP)** | 434 | 171 | 95 | | 45 | | 0.422 | 0.304 | 0.049 | 0.450 |
| **brightarm (EP)** | 434 | | | 170 | | 89 | 0.124 | 0.018 | 0.057 | 0.174 |

*Table A5: Meta information about the models. The first column indicates the number of cases per model. Columns two to six indicate the nested mixed effects structure and the number of mice per grouping levels. Columns seven and eight report the adjusted and unadjusted interclass correlation coefficient. The last two columns indicate the marginal and conditional r-squared. The former corresponds to the variance explained by the fixed effects only and the latter by the entire model including both fixed and random effects.*

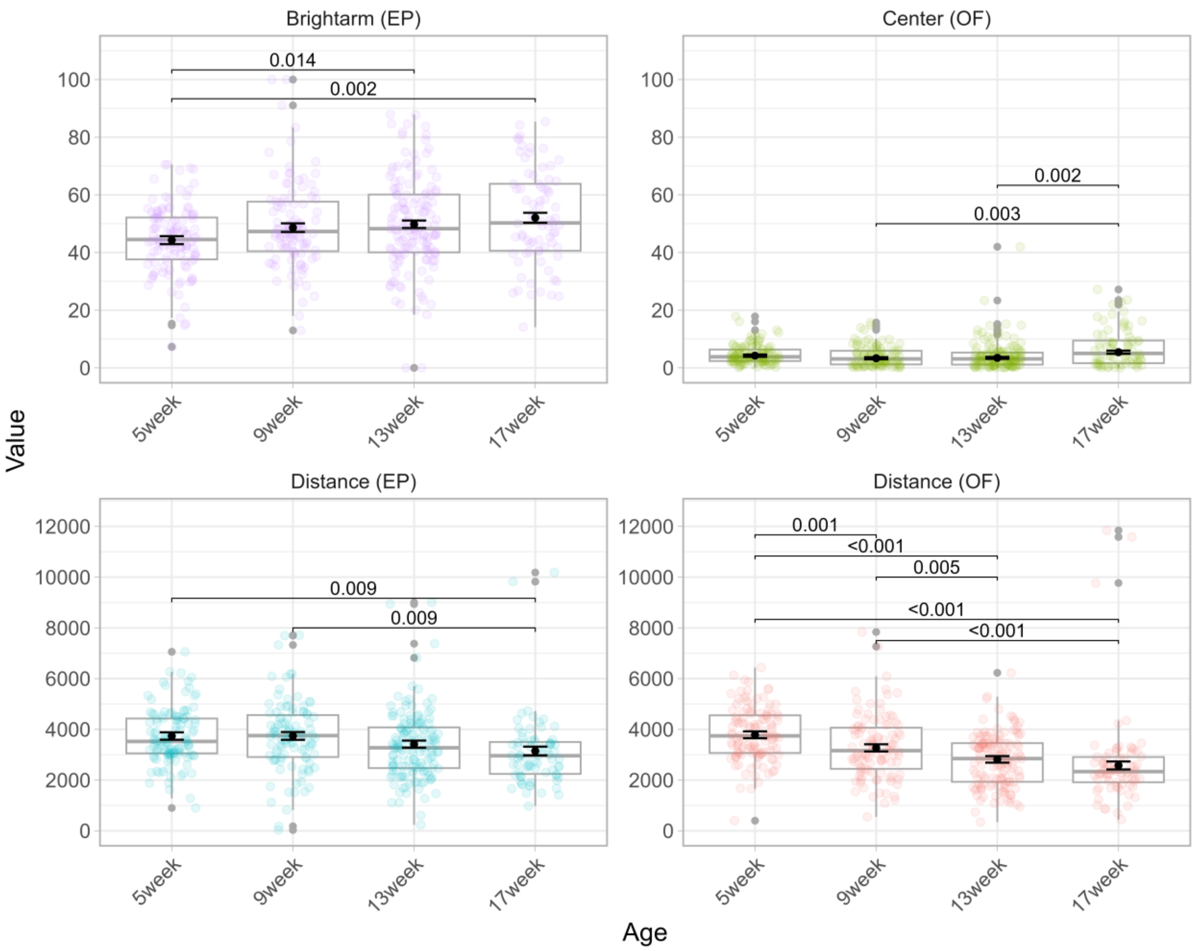


*Figure A3:* Behavioral outcomes across development. Each panel shows boxplots for one behavioral outcome (y-axis) across test ages (x-axis). The black points indicate model-based estimated marginal means for each test age and corresponding standard errors. P-values indicate significant differences between test ages. P-values have been corrected for multiple post-hoc comparisons with the Tukey procedure.

Results in treatment mice: effect of food switch measured at different test ages

#### WEANLINGS

**Table A6. Regression output for all outcome variables (weanlings)**

| | | Kenward-Roger Estimates | | | | | Bootstrapped Estimates | | | |
|---|---|---|---|---|---|---|---|---|---|---|
| | **Starting food** | **Estimate** | ***SE*** | **df** | ***t*** | ***p*** | **Estimate** | **95% CI** | ***p*** | ***p adj.*** |
| **distance (OF)** | | | | | | | | | | |
| sexm | SQ | -81.98 | 160.87 | 41.80 | -0.51 | 0.613 | -80.68 | (-396.048, 227.811) | 0.597 | 0.838 |
| switch_time pregnancy | SQ | 61.69 | 398.67 | 45.43 | 0.15 | 0.878 | 57.46 | (-673.653, 833.272) | 0.875 | 0.875 |
| switch_time birth | SQ | -164.45 | 307.80 | 60.76 | -0.53 | 0.595 | -162.05 | (-774.872, 427.613) | 0.612 | 0.612 |
| sexm | HQ | -56.45 | 149.92 | 141.83 | -0.38 | 0.707 | -54.98 | (-339.972, 232.898) | 0.703 | 0.967 |
| switch_time pregnancy | HQ | -714.07 | 191.32 | 20.22 | -3.73 | **0.001** | -706.23 | **(-1088.111, -350.694)** | **<0.001** | **<0.001** |
| switch_time birth | HQ | -413.23 | 236.45 | 36.26 | -1.75 | 0.089 | -403.33 | (-861.552, 52.786) | 0.081 | 0.162 |
| **center (OF)*** | | | | | | | | | | |
| sexm | SQ* | -0.07 | 0.16 | | -0.42 | 0.675 | -0.07 | (-0.4, 0.26) | 0.666 | 0.838 |
| switch_time pregnancy | SQ* | 0.07 | 0.18 | | 0.37 | 0.711 | 0.07 | (-0.31, 0.47) | 0.740 | 0.875 |
| switch_time birth | SQ* | 0.24 | 0.22 | | 1.10 | 0.274 | 0.23 | (-0.16, 0.66) | 0.244 | 0.558 |
| sexm | HQ* | 0.03 | 0.13 | | 0.25 | 0.807 | 0.04 | (-0.21, 0.29) | 0.796 | 0.967 |
| switch_time pregnancy | HQ* | 0.17 | 0.14 | | 1.18 | 0.238 | 0.17 | (-0.11, 0.43) | 0.249 | 0.332 |
| switch_time birth | HQ* | 0.16 | 0.18 | | 0.85 | 0.399 | 0.15 | (-0.22, 0.56) | 0.436 | 0.444 |
| **distance (EP)** | | | | | | | | | | |
| sexm | SQ | 41.01 | 197.66 | 51.59 | 0.21 | 0.836 | 38.75 | (-338.012, 417.536) | 0.838 | 0.838 |
| switch_time pregnancy | SQ | -187.87 | 286.32 | 13.84 | -0.66 | 0.522 | -198.67 | (-733.376, 361.54) | 0.478 | 0.875 |
| switch_time birth | SQ | 233.49 | 287.27 | 48.43 | 0.81 | 0.420 | 225.90 | (-331.625, 773.489) | 0.410 | 0.558 |
| sexm | HQ | -11.78 | 168.27 | 145.99 | -0.07 | 0.944 | -7.50 | (-339.649, 311.437) | 0.967 | 0.967 |
| switch_time pregnancy | HQ | -308.67 | 267.46 | 22.38 | -1.15 | 0.261 | -311.89 | (-847.78, 207.175) | 0.241 | 0.332 |
| switch_time birth | HQ | -649.55 | 305.93 | 52.33 | -2.12 | **0.038** | -653.03 | **(-1235.237, -72.068)** | **0.027** | 0.109 |
| **brightarm (EP)** | | | | | | | | | | |
| sexm | SQ | -1.14 | 2.39 | 55.02 | -0.47 | 0.637 | -1.10 | (-5.889, 3.563) | 0.640 | 0.838 |
| switch_time pregnancy | SQ | -3.89 | 2.69 | 55.50 | -1.44 | 0.155 | -3.88 | (-9.235, 1.417) | 0.141 | 0.563 |
| switch_time birth | SQ | -2.58 | 3.16 | 55.10 | -0.82 | 0.418 | -2.54 | (-8.695, 3.749) | 0.418 | 0.558 |
| sexm | HQ | -1.09 | 2.27 | 74.65 | -0.48 | 0.632 | -1.08 | (-5.484, 3.171) | 0.635 | 0.967 |
| switch_time pregnancy | HQ | 0.49 | 2.60 | 18.99 | 0.19 | 0.852 | 0.50 | (-4.524, 5.611) | 0.845 | 0.845 |
| switch_time birth | HQ | 2.47 | 3.32 | 30.45 | 0.74 | 0.462 | 2.36 | (-3.864, 8.835) | 0.444 | 0.444 |

*Table A6: Model summary for all outcome variables measured in weanlings. Each combination of outcome variable and starting food corresponds to one model (i.e., eight models in total). All variables were fitted with mixed effects models if possible. We attempted to fit each model with nested random intercepts with cages nested in families. If the model did not converge we simplified the random effects structure (see Table A18). If the model did not converge due to lacking variance from the random effects, we fitted a generalized linear model (GLM) only containing fixed effects (indicated with a *). We applied a square root transformation to Center (OF) because the distribution of times spent in the center was skewed towards the right. For all mixed models degrees of freedom were computed with the Kenward-Roger approximation. Additionally, we computed bootstrapped parameters estimates, confidence intervals, and p-values. We applied a correction for multiple testing (FDR correction) to these bootstrapped p-values across the four outcome variables. All categorical predictors were dummy coded with the following reference groups: females (for sex) and never (for switch time). Center (OF) and brightarm (EP) are measured in percent.*

**Table A7. Anova summary (weanlings)**

| | Starting food | Sum Sq | Mean Sq | NumDF | DenDF | *F* | *p* | *p adj.* |
|---|---|---|---|---|---|---|---|---|
| **distance (OF)** | | | | | | | | |
| sex | SQ | 126151.49 | 126151 | 1.00 | 41.80 | 0.26 | 0.613 | 0.836 |
| switch_time | SQ | 236271.28 | 118136 | 2.00 | 55.48 | 0.24 | 0.785 | 0.785 |
| sex | HQ | 106276.37 | 106276 | 1.00 | 141.83 | 0.14 | 0.707 | 0.944 |
| switch_time | HQ | 10649388.84 | 5324694 | 2.00 | 23.68 | 7.09 | **0.004** | **0.015** |
| **center (OF)*** | | | | | | | | |
| sex | SQ* | 0.14 | | 1.00 | | 0.18 | 0.675 | 0.836 |
| switch_time | SQ* | 0.93 | | 2.00 | | 0.60 | 0.548 | 0.731 |
| sex | HQ* | 0.04 | | 1.00 | | 0.06 | 0.807 | 0.944 |
| switch_time | HQ* | 1.02 | | 2.00 | | 0.79 | 0.454 | 0.605 |
| **distance (EP)** | | | | | | | | |
| sex | SQ | 28357.20 | 28357 | 1.00 | 51.59 | 0.04 | 0.836 | 0.836 |
| switch_time | SQ | 1065221.60 | 532611 | 2.00 | 23.21 | 0.80 | 0.461 | 0.731 |
| sex | HQ | 4231.22 | 4231 | 1.00 | 145.99 | 0.00 | 0.944 | 0.944 |
| switch_time | HQ | 4181615.82 | 2090808 | 2.00 | 29.80 | 2.42 | 0.106 | 0.213 |
| **brightarm (EP)** | | | | | | | | |
| sex | SQ | 26.70 | 27 | 1.00 | 55.02 | 0.22 | 0.637 | 0.836 |
| switch_time | SQ | 259.89 | 130 | 2.00 | 54.90 | 1.09 | 0.342 | 0.731 |
| sex | HQ | 33.45 | 33 | 1.00 | 74.65 | 0.23 | 0.632 | 0.944 |
| switch_time | HQ | 81.52 | 41 | 2.00 | 20.81 | 0.28 | 0.758 | 0.758 |

*Table A7: Analysis of variance summary with Kenward-Roger degrees of freedom. Each combination of outcome variable and starting food corresponds to one model. Per model, we applied a correction for multiple testing (FDR correction) across the four outcome variables which use the same set of predictors. Star symbols (*) indicate models for which it was not possible to estimate a random effects structure. In those cases we estimated a generalized linear model (GLM). Center (OF) and brightarm (EP) are measured in percent.*

**Table A8. Estimated marginal means and medians (weanlings)**

| | | Marginal means | | | | Bootstrapped marginal medians | |
|---|---|---|---|---|---|---|---|
| | Starting food | Mean | *SE* | df | 95% CI | Median | 95% CI |
| **distance (OF)** | | | | | | | |
| never | SQ | 3468.02 | 210.67 | 32.90 | (3039.374, 3896.673) | 3460.95 | (3056.037, 3864.409) |
| pregnancy | SQ | 3529.71 | 351.39 | 36.80 | (2817.604, 4241.825) | 3533.03 | (2850.384, 4206.32) |
| birth | SQ | 3303.57 | 280.30 | 50.27 | (2740.638, 3866.505) | 3300.78 | (2769.544, 3836.897) |
| never | HQ | 4094.59 | 126.08 | 35.77 | (3838.832, 4350.346) | 4091.95 | (3849.945, 4343.81) |
| pregnancy | HQ | 3380.52 | 143.87 | 14.13 | (3072.223, 3688.825) | 3384.41 | (3104.296, 3664.226) |
| birth | HQ | 3681.36 | 203.70 | 25.33 | (3262.102, 4100.609) | 3681.51 | (3288.332, 4078.631) |
| **center (OF)** | | | | | | | |

**Table A8. Estimated marginal means and medians (weanlings)**

| | Starting food | Marginal means | | | | Bootstrapped marginal medians | |
|---|---|---|---|---|---|---|---|
| | | **Mean** | ***SE*** | **df** | **95% CI** | **Median** | **95% CI** |
| never | SQ | 3.88 | 0.47 | 114.00 | (2.948, 4.805) | 3.89 | (3.16, 4.691) |
| pregnancy | SQ | 4.15 | 0.57 | 114.00 | (3.014, 5.287) | 4.16 | (2.957, 5.637) |
| birth | SQ | 4.88 | 0.81 | 114.00 | (3.269, 6.484) | 4.86 | (3.482, 6.667) |
| never | HQ | 4.40 | 0.42 | 146.00 | (3.568, 5.227) | 4.41 | (3.714, 5.219) |
| pregnancy | HQ | 5.14 | 0.47 | 146.00 | (4.21, 6.079) | 5.14 | (4.244, 6.111) |
| birth | HQ | 5.07 | 0.69 | 146.00 | (3.701, 6.445) | 5.08 | (3.716, 6.848) |
| **distance (EP)** | | | | | | | |
| never | SQ | 3518.66 | 165.10 | 31.17 | (3182.017, 3855.312) | 3523.30 | (3195.148, 3830.088) |
| pregnancy | SQ | 3330.79 | 233.58 | 9.41 | (2805.857, 3855.728) | 3342.29 | (2879.502, 3784.206) |
| birth | SQ | 3752.15 | 242.58 | 42.42 | (3262.746, 4241.557) | 3749.09 | (3281.291, 4207.281) |
| never | HQ | 3812.91 | 165.77 | 35.20 | (3476.447, 4149.369) | 3814.54 | (3484.145, 4132.229) |
| pregnancy | HQ | 3504.24 | 209.85 | 17.54 | (3062.527, 3945.958) | 3507.43 | (3078.708, 3923.459) |
| birth | HQ | 3163.36 | 274.31 | 34.84 | (2606.39, 3720.333) | 3158.17 | (2633.124, 3684.079) |
| **brightarm (EP)** | | | | | | | |
| never | SQ | 44.76 | 1.75 | 57.51 | (41.266, 48.261) | 44.74 | (41.324, 48.237) |
| pregnancy | SQ | 40.87 | 2.06 | 54.30 | (36.749, 45) | 40.88 | (36.786, 44.951) |
| birth | SQ | 42.18 | 2.65 | 53.60 | (36.867, 47.497) | 42.11 | (37.063, 47.232) |
| never | HQ | 44.17 | 1.78 | 35.06 | (40.564, 47.78) | 44.19 | (40.805, 47.566) |
| pregnancy | HQ | 44.66 | 1.90 | 12.11 | (40.527, 48.802) | 44.73 | (40.917, 48.369) |
| birth | HQ | 46.64 | 2.80 | 21.49 | (40.827, 52.462) | 46.56 | (41.019, 52.085) |

*Table A8: Estimated marginal means for all categorical predictors. The last two columns indicate bootstrapped marginal medians and confidence intervals. Center (OF) and brightarm (EP) are measured in percent.*

#### EARLY ADOLESCENTS

**Table A9. Regression output for all outcome variables (early adolescents)**

| | Starting food | Kenward-Roger Estimates | | | | | Bootstrapped Estimates | | | |
|---|---|---|---|---|---|---|---|---|---|---|
| | | **Estimate** | ***SE*** | **df** | ***t*** | ***p*** | **Estimate** | **95% CI** | ***p*** | ***p adj.*** |
| **distance (OF)** | | | | | | | | | | |
| sexm | SQ | -137.08 | 180.09 | 60.22 | -0.76 | 0.450 | -136.90 | (-494.485, 224.671) | 0.433 | 0.577 |
| switch_time pregnancy | SQ | 115.78 | 338.08 | 25.01 | 0.34 | 0.735 | 112.59 | (-543.494, 740.296) | 0.740 | 0.740 |
| switch_time birth | SQ | 360.63 | 307.37 | 62.26 | 1.17 | 0.245 | 353.42 | (-232.368, 960.267) | 0.234 | 0.469 |
| switch_time weanling | SQ | 233.34 | 275.55 | 78.89 | 0.85 | 0.400 | 230.39 | (-314.206, 778.197) | 0.404 | 0.452 |
| sexm | HQ | -167.72 | 193.12 | 78.54 | -0.87 | 0.388 | -167.52 | (-540.955, 205.634) | 0.382 | 0.910 |
| switch_time pregnancy | HQ | -616.61 | 270.57 | 25.04 | -2.28 | **0.031** | -618.77 | **(-1142.085, -90.985)** | **0.021** | **0.042** |
| switch_time birth | HQ | -692.19 | 334.05 | 42.76 | -2.07 | **0.044** | -691.48 | **(-1334.853, -43.701)** | **0.037** | 0.147 |
| switch_time weanling | HQ | -143.79 | 307.00 | 91.30 | -0.47 | 0.641 | -140.16 | (-745.14, 449.417) | 0.656 | 0.848 |
| **center (OF)** | | | | | | | | | | |
| sexm | SQ | 0.18 | 0.17 | 64.92 | 1.06 | 0.294 | 0.18 | (-0.16, 0.51) | 0.294 | 0.577 |
| switch_time pregnancy | SQ | 0.42 | 0.22 | 64.44 | 1.95 | 0.055 | 0.42 | (-0.01, 0.84) | 0.054 | 0.214 |

**Table A9. Regression output for all outcome variables (early adolescents)**

| | | Kenward-Roger Estimates | | | | | Bootstrapped Estimates | | | |
|---|---|---|---|---|---|---|---|---|---|---|
| | **Starting food** | **Estimate** | ***SE*** | **df** | ***t*** | ***p*** | **Estimate** | **95% CI** | ***p*** | ***p adj.*** |
| switch_time birth | SQ | 0.49 | 0.26 | 63.01 | 1.92 | 0.060 | 0.50 | (-0.01, 1.01) | 0.057 | 0.227 |
| switch_time weanling | SQ | 0.78 | 0.24 | 66.66 | 3.19 | **0.002** | 0.79 | **(0.31, 1.25)** | **<0.001** | **0.003** |
| sexm | HQ | 0.06 | 0.16 | 78.41 | 0.38 | 0.705 | 0.06 | (-0.25, 0.4) | 0.701 | 0.910 |
| switch_time pregnancy | HQ | 0.51 | 0.19 | 77.84 | 2.63 | **0.010** | 0.51 | **(0.14, 0.88)** | **0.007** | **0.029** |
| switch_time birth | HQ | 0.13 | 0.25 | 79.89 | 0.51 | 0.608 | 0.13 | (-0.35, 0.61) | 0.602 | 0.602 |
| switch_time weanling | HQ | 0.04 | 0.25 | 77.80 | 0.17 | 0.867 | 0.05 | (-0.46, 0.53) | 0.844 | 0.848 |
| **distance (EP)** | | | | | | | | | | |
| sexm | SQ | -528.61 | 208.25 | 118.71 | -2.54 | **0.012** | -527.45 | **(-941.992, -112.589)** | **0.014** | 0.054 |
| switch_time pregnancy | SQ | 242.01 | 384.41 | 26.19 | 0.63 | 0.534 | 232.43 | (-497.788, 966.346) | 0.531 | 0.740 |
| switch_time birth | SQ | 264.32 | 341.05 | 77.17 | 0.78 | 0.441 | 258.60 | (-395.553, 906.816) | 0.418 | 0.557 |
| switch_time weanling | SQ | -226.11 | 309.76 | 109.53 | -0.73 | 0.467 | -229.61 | (-828.488, 378.819) | 0.452 | 0.452 |
| sexm | HQ | -63.27 | 208.43 | 80.32 | -0.30 | 0.762 | -66.88 | (-454.613, 339.559) | 0.751 | 0.910 |
| switch_time pregnancy | HQ | -493.09 | 290.99 | 25.23 | -1.69 | 0.102 | -491.30 | (-1069.999, 83.771) | 0.090 | 0.121 |
| switch_time birth | HQ | -214.79 | 357.53 | 42.24 | -0.60 | 0.551 | -206.17 | (-923.114, 460.51) | 0.550 | 0.602 |
| switch_time weanling | HQ | 59.07 | 327.94 | 89.75 | 0.18 | 0.857 | 63.67 | (-572.593, 688.846) | 0.848 | 0.848 |
| **brightarm (EP)** | | | | | | | | | | |
| sexm | SQ | 1.47 | 2.84 | 119.99 | 0.52 | 0.605 | 1.48 | (-3.962, 6.937) | 0.593 | 0.593 |
| switch_time pregnancy | SQ | -1.74 | 4.47 | 18.19 | -0.39 | 0.701 | -1.83 | (-10.27, 6.808) | 0.665 | 0.740 |
| switch_time birth | SQ | 1.89 | 4.36 | 70.80 | 0.43 | 0.666 | 1.93 | (-6.653, 10.334) | 0.638 | 0.638 |
| switch_time weanling | SQ | -3.47 | 4.08 | 95.68 | -0.85 | 0.398 | -3.48 | (-11.255, 4.27) | 0.378 | 0.452 |
| sexm | HQ | 0.25 | 2.39 | 77.40 | 0.11 | 0.916 | 0.27 | (-4.373, 4.836) | 0.910 | 0.910 |
| switch_time pregnancy | HQ | -0.21 | 3.83 | 25.64 | -0.06 | 0.956 | -0.15 | (-7.467, 7.147) | 0.971 | 0.971 |
| switch_time birth | HQ | -6.16 | 4.52 | 46.17 | -1.36 | 0.180 | -6.05 | (-15.238, 2.735) | 0.168 | 0.335 |
| switch_time weanling | HQ | -4.61 | 3.81 | 87.10 | -1.21 | 0.229 | -4.63 | (-12.031, 2.54) | 0.210 | 0.842 |

*Table A9: Model summary for all outcome variables measured in early adolescents. Each combination of outcome variable and starting food corresponds to one model. All variables were fitted with mixed effects models if possible. We attempted to fit each model with nested random intercepts with cages nested in families. If the model did not converge we simplified the random effects structure (see Table 19). If the model did not converge due to lacking variance from the random effects, we fitted a normal GLM only containing fixed effects (indicated with a *). We applied a square root transformation to Center (OF) because the distribution of times spent in the center was skewed towards the right. For all mixed models degrees of freedom were computed with the Kenward-Roger approximation. Additionally, we computed bootstrapped parameters estimates, confidence intervals, and p-values. We applied a correction for multiple testing (FDR correction) to these bootstrapped p-values across the four outcome variables. All categorical predictors were dummy coded with the following reference groups: females (for sex) and never (for switch time). Center (OF) and brightarm (EP) are measured in percent.*

**Table A10. Anova summary (early adolescents)**

| | **Starting food** | **Sum Sq** | **Mean Sq** | **NumDF** | **DenDF** | ***F*** | ***p*** | ***p adj.*** |
|---|---|---|---|---|---|---|---|---|
| **distance (OF)** | | | | | | | | |
| sex | SQ | 418171.09 | 418171 | 1 | 60.22 | 0.58 | 0.450 | 0.599 |
| switch_time | SQ | 1254994.69 | 418332 | 3 | 46.54 | 0.57 | 0.634 | 0.717 |
| sex | HQ | 790926.12 | 790926 | 1 | 78.54 | 0.75 | 0.388 | 0.916 |
| switch_time | HQ | 7593299.08 | 2531100 | 3 | 37.39 | 2.39 | 0.084 | 0.168 |
| **center (OF)** | | | | | | | | |
| sex | SQ | 0.99 | 1 | 1 | 64.92 | 1.12 | 0.294 | 0.588 |
| switch_time | SQ | 10.03 | 3 | 3 | 64.39 | 3.77 | **0.015** | 0.059 |

| Table A10. Anova summary (early adolescents) | | | | | | | | |
|---|---|---|---|---|---|---|---|---|
| | **Starting food** | **Sum Sq** | **Mean Sq** | **NumDF** | **DenDF** | ***F*** | ***p*** | ***p adj.*** |
| sex | HQ | 0.12 | 0 | 1 | 78.41 | 0.14 | 0.705 | 0.916 |
| switch_time | HQ | 6.73 | 2 | 3 | 78.66 | 2.66 | 0.054 | 0.168 |
| **distance (EP)** | | | | | | | | |
| sex | SQ | 7445096.77 | 7445097 | 1 | 118.71 | 6.44 | **0.012** | 0.050 |
| switch_time | SQ | 2197674.10 | 732558 | 3 | 51.40 | 0.63 | 0.599 | 0.717 |
| sex | HQ | 107124.12 | 107124 | 1 | 80.32 | 0.09 | 0.762 | 0.916 |
| switch_time | HQ | 3985089.09 | 1328363 | 3 | 37.26 | 1.13 | 0.348 | 0.381 |
| **brightarm (EP)** | | | | | | | | |
| sex | SQ | 61.42 | 61 | 1 | 119.99 | 0.27 | 0.605 | 0.605 |
| switch_time | SQ | 311.85 | 104 | 3 | 37.97 | 0.45 | 0.717 | 0.717 |
| sex | HQ | 1.77 | 2 | 1 | 77.40 | 0.01 | 0.916 | 0.916 |
| switch_time | HQ | 500.54 | 167 | 3 | 41.51 | 1.05 | 0.381 | 0.381 |

*Table A10: Analysis of variance summary with Kenward-Roger degrees of freedom. Each combination of outcome variable and starting food corresponds to one model. Per model, we applied a correction for multiple testing (FDR correction) across the four outcome variables which use the same set of predictors. Star symbols (*) indicate models for which it was not possible to estimate a random effects structure. In those cases we estimated a generalized linear model (GLM). Center (OF) and brightarm (EP) are measured in percent.*

| Table A11. Estimated marginal means and medians (early adolescents) | | | | | | | |
|---|---|---|---|---|---|---|---|
| | | **Marginal means** | | | | **Bootstrapped marginal medians** | |
| | **Starting food** | **Mean** | ***SE*** | **df** | **95% CI** | **Median** | **95% CI** |
| **distance (OF)** | | | | | | | |
| never | SQ | 2799.09 | 189.53 | 40.15 | (2416.077, 3182.108) | 2791.86 | (2424.522, 3172.791) |
| pregnancy | SQ | 2914.87 | 280.47 | 18.47 | (2326.711, 3503.033) | 2911.78 | (2350.484, 3435.068) |
| birth | SQ | 3159.72 | 263.50 | 53.61 | (2631.353, 3688.094) | 3155.49 | (2654.293, 3688.575) |
| weanling | SQ | 3032.43 | 243.00 | 52.99 | (2545.034, 3519.832) | 3036.32 | (2546.366, 3505.056) |
| never | HQ | 3688.52 | 182.40 | 46.13 | (3321.404, 4055.645) | 3694.55 | (3337.924, 4051.264) |
| pregnancy | HQ | 3071.92 | 199.84 | 16.03 | (2648.328, 3495.507) | 3074.51 | (2688.902, 3472.237) |
| birth | HQ | 2996.33 | 285.95 | 32.06 | (2413.915, 3578.747) | 2995.35 | (2448.856, 3524.751) |
| weanling | HQ | 3544.73 | 276.65 | 62.26 | (2991.769, 4097.693) | 3538.00 | (3026.977, 4071.254) |
| **center (OF)** | | | | | | | |
| never | SQ | 3.03 | 0.50 | 63.50 | (2.031, 4.022) | 3.02 | (2.131, 4.056) |
| pregnancy | SQ | 4.66 | 0.69 | 62.82 | (3.279, 6.048) | 4.67 | (3.447, 6.124) |
| birth | SQ | 4.98 | 0.95 | 61.97 | (3.077, 6.891) | 5.00 | (3.33, 7.066) |
| weanling | SQ | 6.35 | 1.00 | 64.72 | (4.363, 8.342) | 6.37 | (4.595, 8.459) |
| never | HQ | 3.43 | 0.52 | 77.91 | (2.402, 4.461) | 3.44 | (2.501, 4.527) |
| pregnancy | HQ | 5.58 | 0.64 | 77.93 | (4.313, 6.849) | 5.58 | (4.458, 6.904) |
| birth | HQ | 3.92 | 0.83 | 79.75 | (2.279, 5.57) | 3.95 | (2.475, 5.756) |
| weanling | HQ | 3.59 | 0.81 | 77.82 | (1.979, 5.204) | 3.63 | (2.167, 5.328) |
| **distance (EP)** | | | | | | | |
| never | SQ | 3443.55 | 215.10 | 38.71 | (3008.367, 3878.735) | 3448.16 | (3020.376, 3861.347) |
| pregnancy | SQ | 3685.56 | 319.41 | 20.13 | (3019.554, 4351.569) | 3682.68 | (3074.059, 4302.505) |
| birth | SQ | 3707.87 | 289.75 | 56.81 | (3127.606, 4288.133) | 3706.89 | (3150.539, 4270.605) |
| weanling | SQ | 3217.44 | 270.04 | 51.42 | (2675.417, 3759.46) | 3207.14 | (2678.485, 3726.426) |
| never | HQ | 3818.34 | 196.05 | 46.14 | (3423.741, 4212.94) | 3821.76 | (3434.751, 4206.901) |

**Table A11. Estimated marginal means and medians (early adolescents)**

| | Starting food | Marginal means Mean | *SE* | df | 95% CI | Bootstrapped marginal medians Median | 95% CI |
|---|---|---|---|---|---|---|---|
| pregnancy | HQ | 3325.25 | 215.32 | 16.37 | (2869.633, 3780.867) | 3326.47 | (2901.307, 3755.251) |
| birth | HQ | 3603.55 | 305.70 | 31.46 | (2980.453, 4226.657) | 3610.38 | (3014.805, 4203.479) |
| weanling | HQ | 3877.41 | 295.31 | 61.01 | (3286.9, 4467.929) | 3887.81 | (3310.198, 4439.464) |
| **brightarm (EP)** | | | | | | | |
| never | SQ | 47.73 | 2.62 | 34.81 | (42.406, 53.063) | 47.70 | (42.584, 52.946) |
| pregnancy | SQ | 45.99 | 3.61 | 13.07 | (38.205, 53.784) | 46.04 | (39.136, 53.161) |
| birth | SQ | 49.63 | 3.63 | 55.63 | (42.353, 56.899) | 49.50 | (42.491, 56.594) |
| weanling | SQ | 44.27 | 3.36 | 51.77 | (37.527, 51.003) | 44.22 | (37.678, 50.811) |
| never | HQ | 50.70 | 2.46 | 43.65 | (45.741, 55.661) | 50.69 | (45.904, 55.431) |
| pregnancy | HQ | 50.49 | 2.93 | 18.39 | (44.334, 56.641) | 50.44 | (44.711, 56.199) |
| birth | HQ | 44.54 | 3.95 | 34.69 | (36.519, 52.571) | 44.61 | (37.024, 51.999) |
| weanling | HQ | 46.09 | 3.62 | 66.39 | (38.87, 53.307) | 46.11 | (39.08, 53.125) |

*Table A11: Estimated marginal means for all categorical predictors. The last two columns indicate bootstrapped marginal medians and confidence intervals. Center (OF) and brightarm (EP) are measured in percent.*

### LATE ADOLESCENTS

**Table A12. Regression output for all outcome variables (late adolescents)**

| | Starting food | Kenward-Roger Estimates Estimate | *SE* | df | *t* | *p* | Bootstrapped Estimates Estimate | 95% CI | *p* | *p adj.* |
|---|---|---|---|---|---|---|---|---|---|---|
| **distance (OF)** | | | | | | | | | | |
| sexm | SQ | 83.29 | 151.24 | 127.12 | 0.55 | 0.583 | 83.61 | (-208.597, 380.67) | 0.572 | 0.763 |
| switch_time pregnancy | SQ | -108.04 | 351.52 | 20.47 | -0.31 | 0.762 | -118.01 | (-784.92, 570.899) | 0.727 | 0.727 |
| switch_time weanling | SQ | 213.90 | 227.05 | 129.64 | 0.94 | 0.348 | 211.75 | (-242.789, 650.011) | 0.359 | 0.741 |
| sexm | HQ | -128.44 | 175.16 | 69.42 | -0.73 | 0.466 | -131.28 | (-472.325, 207.836) | 0.468 | 0.623 |
| switch_time pregnancy | HQ | -636.53 | 225.50 | 21.43 | -2.82 | **0.010** | -633.98 | **(-1053.94, -189.762)** | **0.003** | **0.006** |
| switch_time weanling | HQ | 23.56 | 272.41 | 82.83 | 0.09 | 0.931 | 25.13 | (-493.589, 551.409) | 0.932 | 0.932 |
| **center (OF)** | | | | | | | | | | |
| sexm | SQ | 0.11 | 0.19 | 51.94 | 0.58 | 0.565 | 0.11 | (-0.26, 0.47) | 0.548 | 0.763 |
| switch_time pregnancy | SQ | 0.48 | 0.22 | 9.23 | 2.15 | 0.059 | 0.48 | **(0.06, 0.88)** | **0.023** | 0.091 |
| switch_time weanling | SQ | 0.15 | 0.26 | 67.00 | 0.56 | 0.577 | 0.15 | (-0.34, 0.62) | 0.556 | 0.741 |
| sexm | HQ | 0.08 | 0.18 | 66.39 | 0.44 | 0.658 | 0.08 | (-0.27, 0.44) | 0.660 | 0.660 |
| switch_time pregnancy | HQ | 0.34 | 0.19 | 65.19 | 1.81 | 0.075 | 0.34 | (-0.03, 0.73) | 0.071 | 0.095 |
| switch_time weanling | HQ | -0.20 | 0.27 | 69.88 | -0.74 | 0.464 | -0.20 | (-0.74, 0.33) | 0.462 | 0.772 |
| **distance (EP)** | | | | | | | | | | |
| sexm | SQ | -160.07 | 244.02 | 53.51 | -0.66 | 0.515 | -158.34 | (-613.932, 314.786) | 0.513 | 0.763 |
| switch_time pregnancy | SQ | -252.59 | 329.61 | 12.43 | -0.77 | 0.458 | -257.53 | (-895.157, 390.039) | 0.438 | 0.583 |
| switch_time weanling | SQ | 42.02 | 341.60 | 69.77 | 0.12 | 0.902 | 35.76 | (-601.451, 692.813) | 0.910 | 0.910 |
| sexm | HQ | -152.72 | 201.72 | 64.94 | -0.76 | 0.452 | -149.80 | (-548.772, 232.463) | 0.436 | 0.623 |
| switch_time pregnancy | HQ | -738.73 | 216.89 | 64.27 | -3.41 | **0.001** | -732.73 | **(-1151.554, -318.524)** | **0.001** | **0.005** |
| switch_time weanling | HQ | 163.43 | 300.55 | 68.23 | 0.54 | 0.588 | 166.31 | (-416.711, 748.54) | 0.579 | 0.772 |

**Table A12. Regression output for all outcome variables (late adolescents)**

| | | Kenward-Roger Estimates | | | | | Bootstrapped Estimates | | | |
|---|---|---|---|---|---|---|---|---|---|---|
| | **Starting food** | **Estimate** | ***SE*** | **df** | ***t*** | ***p*** | **Estimate** | **95% CI** | ***p*** | ***p adj.*** |
| **brightarm (EP)** | | | | | | | | | | |
| sexm | SQ | 0.22 | 3.30 | 51.58 | 0.07 | 0.947 | 0.08 | (-6.199, 6.639) | 0.978 | 0.978 |
| switch_time pregnancy | SQ | -4.24 | 4.08 | 10.40 | -1.04 | 0.323 | -4.21 | (-11.92, 3.477) | 0.293 | 0.583 |
| switch_time weanling | SQ | -3.31 | 4.63 | 66.06 | -0.71 | 0.478 | -3.17 | (-11.884, 5.698) | 0.477 | 0.741 |
| sexm | HQ | -2.96 | 2.61 | 65.31 | -1.14 | 0.260 | -3.02 | (-8.168, 2.208) | 0.255 | 0.623 |
| switch_time pregnancy | HQ | -0.73 | 2.81 | 64.57 | -0.26 | 0.796 | -0.73 | (-6.033, 4.834) | 0.800 | 0.800 |
| switch_time weanling | HQ | 2.95 | 3.89 | 68.13 | 0.76 | 0.451 | 2.98 | (-4.914, 10.443) | 0.439 | 0.772 |

*Table A12: Model summary for all outcome variables measured in late adolescents. Each combination of outcome variable and starting food corresponds to one model. All variables were fitted with mixed effects models if possible. We attempted to fit each model with nested random intercepts with cages nested in families. If the model did not converge we simplified the random effects structure (see Table 19). If the model did not converge due to lacking variance from the random effects, we fitted a normal GLM only containing fixed effects (indicated with a *). We applied a square root transformation to Center (OF) because the distribution of times spent in the center was skewed towards the right. For all mixed models degrees of freedom were computed with the Kenward-Roger approximation. Additionally, we computed bootstrapped parameters estimates, confidence intervals, and p-values. We applied a correction for multiple testing (FDR correction) to these bootstrapped p-values across the four outcome variables. All categorical predictors were dummy coded with the following reference groups: females (for sex) and never (for switch time). Center (OF) and brightarm (EP) are measured in percent.*

**Table A13. Anova summary (late adolescents)**

| | **Starting food** | **Sum Sq** | **Mean Sq** | **NumDF** | **DenDF** | ***F*** | ***p*** | ***p adj.*** |
|---|---|---|---|---|---|---|---|---|
| **distance (OF)** | | | | | | | | |
| sex | SQ | 170316.55 | 170317 | 1 | 127.12 | 0.30 | 0.583 | 0.777 |
| switch_time | SQ | 624741.43 | 312371 | 2 | 40.76 | 0.55 | 0.581 | 0.708 |
| sex | HQ | 421805.32 | 421805 | 1 | 69.42 | 0.54 | 0.466 | 0.621 |
| switch_time | HQ | 6743048.14 | 3371524 | 2 | 34.85 | 4.25 | **0.022** | **0.045** |
| **center (OF)** | | | | | | | | |
| sex | SQ | 0.33 | 0 | 1 | 51.94 | 0.34 | 0.565 | 0.777 |
| switch_time | SQ | 4.59 | 2 | 2 | 16.36 | 2.26 | 0.136 | 0.543 |
| sex | HQ | 0.17 | 0 | 1 | 66.39 | 0.20 | 0.658 | 0.658 |
| switch_time | HQ | 4.64 | 2 | 2 | 68.99 | 2.67 | 0.077 | 0.102 |
| **distance (EP)** | | | | | | | | |
| sex | SQ | 418627.84 | 418628 | 1 | 53.51 | 0.43 | 0.515 | 0.777 |
| switch_time | SQ | 694568.81 | 347284 | 2 | 21.71 | 0.35 | 0.708 | 0.708 |
| sex | HQ | 723277.41 | 723277 | 1 | 64.94 | 0.57 | 0.452 | 0.621 |
| switch_time | HQ | 18596880.50 | 9298440 | 2 | 68.12 | 7.37 | **0.001** | **0.005** |
| **brightarm (EP)** | | | | | | | | |
| sex | SQ | 1.19 | 1 | 1 | 51.58 | 0.00 | 0.947 | 0.947 |
| switch_time | SQ | 334.24 | 167 | 2 | 18.27 | 0.62 | 0.551 | 0.708 |
| sex | HQ | 261.02 | 261 | 1 | 65.31 | 1.29 | 0.260 | 0.621 |
| switch_time | HQ | 175.97 | 88 | 2 | 68.10 | 0.43 | 0.649 | 0.649 |

*Table A13: Analysis of variance summary with Kenward-Roger degrees of freedom. Each combination of outcome variable and starting food corresponds to one model. Per model, we applied a correction for multiple testing (FDR correction) across the four outcome variables which use the same set of predictors. Star symbols (*) indicate models for which it was not possible to estimate a random effects structure. In those cases we estimated a generalized linear model (GLM). Center (OF) and brightarm (EP) are measured in percent.*

**Table A14. Estimated marginal means and medians (late adolescents)**

| | | Marginal means | | | | Bootstrapped marginal medians | |
|---|---|---|---|---|---|---|---|
| | Starting food | Mean | *SE* | df | 95% CI | Median | 95% CI |
| **distance (OF)** | | | | | | | |
| never | SQ | 2603.10 | 168.25 | 27.44 | (2258.14, 2948.056) | 2605.92 | (2285.617, 2943.912) |
| pregnancy | SQ | 2495.05 | 308.82 | 18.87 | (1848.396, 3141.712) | 2504.16 | (1893.477, 3128.603) |
| weanling | SQ | 2817.00 | 228.99 | 61.03 | (2359.106, 3274.891) | 2816.12 | (2368.295, 3257.487) |
| never | HQ | 3070.66 | 142.44 | 40.03 | (2782.784, 3358.536) | 3069.28 | (2790.146, 3355.586) |
| pregnancy | HQ | 2434.13 | 174.88 | 14.71 | (2060.753, 2807.511) | 2437.24 | (2088.511, 2765.518) |
| weanling | HQ | 3094.22 | 254.65 | 62.65 | (2585.293, 3603.143) | 3101.43 | (2601.457, 3579.454) |
| **center (OF)** | | | | | | | |
| never | SQ | 2.95 | 0.45 | 35.58 | (2.041, 3.862) | 2.96 | (2.175, 3.86) |
| pregnancy | SQ | 4.81 | 0.78 | 8.60 | (3.028, 6.59) | 4.82 | (3.467, 6.363) |
| weanling | SQ | 3.47 | 0.82 | 58.70 | (1.828, 5.114) | 3.49 | (2.104, 5.186) |
| never | HQ | 3.73 | 0.51 | 54.19 | (2.714, 4.746) | 3.75 | (2.827, 4.812) |
| pregnancy | HQ | 5.18 | 0.63 | 70.44 | (3.928, 6.433) | 5.18 | (4.032, 6.514) |
| weanling | HQ | 3.01 | 0.81 | 73.81 | (1.4, 4.62) | 3.03 | (1.609, 4.78) |
| **distance (EP)** | | | | | | | |
| never | SQ | 3248.54 | 186.26 | 29.93 | (2868.12, 3628.97) | 3249.08 | (2888.776, 3617.448) |
| pregnancy | SQ | 2995.95 | 272.10 | 8.64 | (2376.439, 3615.467) | 3002.98 | (2476.461, 3529.243) |
| weanling | SQ | 3290.57 | 294.68 | 54.89 | (2699.99, 3881.145) | 3288.14 | (2736.603, 3846.621) |
| never | HQ | 3583.54 | 146.92 | 52.75 | (3288.82, 3878.264) | 3583.53 | (3306.45, 3862.71) |
| pregnancy | HQ | 2844.81 | 159.79 | 76.99 | (2526.62, 3162.997) | 2845.04 | (2533.158, 3159.747) |
| weanling | HQ | 3746.97 | 262.96 | 75.28 | (3223.159, 4270.78) | 3753.48 | (3255.732, 4260.699) |
| **brightarm (EP)** | | | | | | | |
| never | SQ | 52.16 | 2.37 | 26.57 | (47.299, 57.014) | 52.15 | (47.632, 56.702) |
| pregnancy | SQ | 47.92 | 3.33 | 6.77 | (40, 55.84) | 47.95 | (41.505, 54.11) |
| weanling | SQ | 48.85 | 3.97 | 56.75 | (40.899, 56.804) | 48.86 | (41.189, 56.458) |
| never | HQ | 47.77 | 1.91 | 53.87 | (43.95, 51.591) | 47.75 | (44.091, 51.437) |
| pregnancy | HQ | 47.04 | 2.07 | 76.31 | (42.928, 51.156) | 47.03 | (42.93, 51.197) |
| weanling | HQ | 50.72 | 3.40 | 74.64 | (43.945, 57.492) | 50.69 | (44.219, 57.316) |

*Table A14: Estimated marginal means for all categorical predictors. The last two columns indicate bootstrapped marginal medians and confidence intervals. Center (OF) and brightarm (EP) are measured in percent.*

### ADULTS

**Table A15. Regression output for all outcome variables (adults)**

| | | Kenward-Roger Estimates | | | | | Bootstrapped Estimates | | | |
|---|---|---|---|---|---|---|---|---|---|---|
| | Starting food | Estimate | *SE* | df | *t* | *p* | Estimate | 95% CI | *p* | *p adj.* |
| **distance (OF)** | | | | | | | | | | |
| sexm | SQ | -115.96 | 183.18 | 77.71 | -0.63 | 0.529 | -116.84 | (-481.724, 238.49) | 0.532 | 0.702 |
| switch_time adult | SQ | 40.45 | 180.04 | 77.50 | 0.22 | 0.823 | 40.15 | (-306.076, 394.332) | 0.817 | 0.817 |
| sexm | HQ | 496.89 | 197.22 | 35.19 | 2.52 | **0.016** | 500.01 | **(112.695, 883.759)** | **0.011** | **0.045** |
| switch_time adult | HQ | 306.59 | 197.51 | 36.27 | 1.55 | 0.129 | 304.41 | (-79.505, 686.487) | 0.120 | 0.333 |

**Table A15. Regression output for all outcome variables (adults)**

| | Starting food | Kenward-Roger Estimates Estimate | *SE* | df | *t* | *p* | Bootstrapped Estimates Estimate | 95% CI | *p* | *p adj.* |
|---|---|---|---|---|---|---|---|---|---|---|
| **center (OF)*** | | | | | | | | | | |
| sexm | SQ | 0.56 | 0.27 | 70.62 | 2.04 | **0.045** | 0.56 | **(0.04, 1.09)** | **0.036** | 0.144 |
| switch_time adult | SQ | -0.27 | 0.27 | 76.59 | -1.00 | 0.319 | -0.27 | (-0.8, 0.25) | 0.291 | 0.742 |
| sexm | HQ* | 0.18 | 0.23 | | 0.78 | 0.440 | 0.18 | (-0.28, 0.63) | 0.433 | 0.804 |
| switch_time adult | HQ* | -0.23 | 0.23 | | -0.97 | 0.334 | -0.22 | (-0.71, 0.23) | 0.333 | 0.333 |
| **distance (EP)** | | | | | | | | | | |
| sexm | SQ | -289.61 | 282.76 | 42.60 | -1.02 | 0.312 | -288.14 | (-840.08, 249.62) | 0.290 | 0.579 |
| switch_time adult | SQ | 249.42 | 279.92 | 41.36 | 0.89 | 0.378 | 245.03 | (-297.101, 792.502) | 0.371 | 0.742 |
| sexm | HQ | -223.18 | 444.61 | 38.01 | -0.50 | 0.619 | -217.58 | (-1072.318, 605.867) | 0.603 | 0.804 |
| switch_time adult | HQ | -426.02 | 435.15 | 42.66 | -0.98 | 0.333 | -433.32 | (-1240.069, 381.485) | 0.300 | 0.333 |
| **brightarm (EP)*** | | | | | | | | | | |
| sexm | SQ | -1.51 | 3.72 | 39.93 | -0.40 | 0.688 | -1.45 | (-8.929, 5.887) | 0.702 | 0.702 |
| switch_time adult | SQ | -1.68 | 3.73 | 40.49 | -0.45 | 0.655 | -1.64 | (-9.07, 5.821) | 0.659 | 0.817 |
| sexm | HQ* | 0.43 | 4.70 | | 0.09 | 0.927 | 0.45 | (-9.001, 9.491) | 0.918 | 0.918 |
| switch_time adult | HQ* | -6.00 | 4.69 | | -1.28 | 0.205 | -6.11 | (-15.413, 2.894) | 0.184 | 0.333 |

*Table A15: Model summary for all outcome variables measured in adults. Each combination of outcome variable and starting food corresponds to one model. All variables were fitted with mixed effects models if possible. We attempted to fit each model with nested random intercepts with cages nested in families. If the model did not converge we simplified the random effects structure (see Table 19). If the model did not converge due to lacking variance from the random effects, we fitted a normal GLM only containing fixed effects (indicated with a *). We applied a square root transformation to Center (OF) because the distribution of times spent in the center was skewed towards the right. For all mixed models degrees of freedom were computed with the Kenward-Roger approximation. Additionally, we computed bootstrapped parameters estimates, confidence intervals, and p-values. We applied a correction for multiple testing (FDR correction) to these bootstrapped p-values across the four outcome variables. All categorical predictors were dummy coded with the following reference groups: females (for sex) and never (for switch time). Center (OF) and brightarm (EP) are measured in percent.*

**Table A16. Anova summary (adults)**

| | Starting food | Sum Sq | Mean Sq | NumDF | DenDF | *F* | *p* | *p adj.* |
|---|---|---|---|---|---|---|---|---|
| **distance (OF)** | | | | | | | | |
| sex | SQ | 184082.83 | 184083 | 1.00 | 77.71 | 0.40 | 0.529 | 0.688 |
| switch_time | SQ | 23185.32 | 23185 | 1.00 | 77.50 | 0.05 | 0.823 | 0.823 |
| sex | HQ | 4448438.11 | 4448438 | 1.00 | 35.19 | 6.35 | **0.016** | 0.066 |
| switch_time | HQ | 1688661.35 | 1688661 | 1.00 | 36.27 | 2.41 | 0.129 | 0.334 |
| **center (OF)*** | | | | | | | | |
| Sex | SQ | 5.49 | 5 | 1.00 | 70.62 | 4.15 | **0.045** | 0.181 |
| switch_time | SQ | 1.33 | 1 | 1.00 | 76.59 | 1.01 | 0.319 | 0.756 |
| sex | HQ* | 0.65 | | 1.00 | | 0.60 | 0.440 | 0.825 |
| switch_time | HQ* | 1.01 | | 1.00 | | 0.94 | 0.334 | 0.334 |
| **distance (EP)** | | | | | | | | |
| sex | SQ | 540545.89 | 540546 | 1.00 | 42.60 | 1.05 | 0.312 | 0.623 |
| switch_time | SQ | 409089.25 | 409089 | 1.00 | 41.36 | 0.79 | 0.378 | 0.756 |
| sex | HQ | 540608.92 | 540609 | 1.00 | 38.01 | 0.25 | 0.619 | 0.825 |
| switch_time | HQ | 2056428.14 | 2056428 | 1.00 | 42.66 | 0.96 | 0.333 | 0.334 |
| **brightarm (EP)*** | | | | | | | | |
| sex | SQ | 24.94 | 25 | 1.00 | 39.93 | 0.16 | 0.688 | 0.688 |
| switch_time | SQ | 30.91 | 31 | 1.00 | 40.49 | 0.20 | 0.655 | 0.823 |

**Table A16. Anova summary (adults)**

| | Starting food | Sum Sq | Mean Sq | NumDF | DenDF | *F* | *p* | *p adj.* |
|---|---|---|---|---|---|---|---|---|
| sex | HQ* | 3.57 | | 1.00 | | 0.01 | 0.927 | 0.927 |
| switch_time | HQ* | 699.12 | | 1.00 | | 1.63 | 0.205 | 0.334 |

*Table A16: Analysis of variance summary with Kenward-Roger degrees of freedom. Each combination of outcome variable and starting food corresponds to one model. Per model, we applied a correction for multiple testing (FDR correction) across the four outcome variables which use the same set of predictors. Star symbols (*) indicate models for which it was not possible to estimate a random effects structure. In those cases we estimated a generalized linear model (GLM). Center (OF) and brightarm (EP) are measured in percent.*

**Table A17. Estimated marginal means and medians (adults)**

| | | Marginal means | | | | Bootstrapped marginal medians | |
|---|---|---|---|---|---|---|---|
| | **Starting food** | **Mean** | ***SE*** | **df** | **95% CI** | **Median** | **95% CI** |
| **distance (OF)** | | | | | | | |
| never | SQ | 2115.85 | 169.71 | 35.30 | (1771.425, 2460.27) | 2118.82 | (1794.089, 2442.306) |
| adult | SQ | 2156.29 | 157.96 | 28.65 | (1833.055, 2479.533) | 2152.75 | (1848.229, 2463.057) |
| never | HQ | 2569.05 | 143.89 | 36.24 | (2277.287, 2860.806) | 2568.15 | (2289.831, 2858.339) |
| adult | HQ | 2875.64 | 134.89 | 35.01 | (2601.794, 3149.481) | 2875.23 | (2606.288, 3135.448) |
| **center (OF)** | | | | | | | |
| never | SQ | 4.35 | 0.87 | 43.25 | (2.596, 6.108) | 4.39 | (2.876, 6.176) |
| adult | SQ | 3.31 | 0.68 | 33.60 | (1.926, 4.688) | 3.31 | (2.161, 4.764) |
| never | HQ | 6.48 | 0.85 | 76.00 | (4.794, 8.175) | 6.48 | (4.848, 8.447) |
| adult | HQ | 5.38 | 0.76 | 76.00 | (3.87, 6.892) | 5.38 | (4.131, 6.813) |
| **distance (EP)** | | | | | | | |
| never | SQ | 2816.51 | 212.82 | 35.74 | (2384.77, 3248.241) | 2817.91 | (2413.106, 3228.216) |
| adult | SQ | 3065.93 | 190.79 | 27.11 | (2674.535, 3457.316) | 3064.62 | (2701.058, 3431.751) |
| never | HQ | 3263.50 | 310.20 | 32.92 | (2632.337, 3894.669) | 3258.17 | (2675.041, 3862.2) |
| adult | HQ | 2837.48 | 317.04 | 25.33 | (2184.948, 3490.008) | 2839.47 | (2246.943, 3421.685) |
| **brightarm (EP)** | | | | | | | |
| never | SQ | 51.49 | 2.80 | 40.36 | (45.824, 57.155) | 51.47 | (45.961, 57.172) |
| adult | SQ | 49.81 | 2.46 | 39.88 | (44.83, 54.784) | 49.81 | (45, 54.596) |
| never | HQ | 52.94 | 3.32 | 75.00 | (46.319, 59.566) | 52.96 | (47.963, 58.25) |
| adult | HQ | 46.95 | 3.31 | 75.00 | (40.349, 53.544) | 46.94 | (39.396, 54.469) |

*Table A17: Estimated marginal means for all categorical predictors. The last two columns indicate bootstrapped marginal medians and confidence intervals. Center (OF) and brightarm (EP) are measured in percent.*

**Table A18. Meta information models**

| Outcome | starting_food | testage | n | cage_family | family | cage | ICC_adj | ICC_unadj | r2_m | r2_c |
|---|---|---|---|---|---|---|---|---|---|---|
| **distance (OF)** | SQ | weanlings | 118 | 67 | 27 | | 0.583 | 0.349 | 0.007 | 0.586 |
| **distance (OF)** | SQ | earlyAdolescents | 136 | 80 | 27 | | 0.319 | 0.088 | 0.021 | 0.333 |
| **distance (OF)** | SQ | lateAdolescents | 134 | | 25 | | 0.385 | 0.007 | 0.013 | 0.393 |
| **distance (OF)** | SQ | adults | 81 | | 22 | | 0.350 | 0.041 | 0.006 | 0.353 |
| **distance (OF)** | HQ | weanlings | 150 | | 29 | | 0.068 | 0.007 | 0.117 | 0.177 |
| **distance (OF)** | HQ | earlyAdolescents | 166 | 92 | 30 | | 0.213 | 0.066 | 0.072 | 0.269 |
| **distance (OF)** | HQ | lateAdolescents | 149 | 82 | 31 | | 0.198 | 0.034 | 0.098 | 0.276 |
| **distance (OF)** | HQ | adults | 76 | | | 42 | 0.011 | 0.002 | 0.115 | 0.125 |
| **distance (EP)** | SQ | weanlings | 114 | 67 | 27 | | 0.306 | 0.002 | 0.024 | 0.322 |

**Table A18. Meta information models**

| Outcome | starting_food | testage | n | cage_family | family | cage | ICC_adj | ICC_unadj | r2_m | r2_c |
|---|---|---|---|---|---|---|---|---|---|---|
| **distance (EP)** | SQ | earlyAdolescents | 125 | | 26 | | 0.183 | 0.002 | 0.064 | 0.235 |
| **distance (EP)** | SQ | lateAdolescents | 133 | 67 | 25 | | 0.315 | 0.063 | 0.015 | 0.325 |
| **distance (EP)** | SQ | adults | 82 | 47 | 22 | | 0.514 | 0.152 | 0.037 | 0.531 |
| **distance (EP)** | HQ | weanlings | 150 | | 29 | | 0.200 | 0.008 | 0.050 | 0.241 |
| **distance (EP)** | HQ | earlyAdolescents | 163 | 92 | 30 | | 0.227 | 0.058 | 0.036 | 0.255 |
| **distance (EP)** | HQ | lateAdolescents | 146 | | | 73 | 0.063 | 0.000 | 0.103 | 0.160 |
| **distance (EP)** | HQ | adults | 77 | 45 | 25 | | 0.222 | 0.018 | 0.022 | 0.239 |
| **center (OF)*** | SQ | weanlings | 118 | - | - | - | - | - | 0.012 | 0.000 |
| **center (OF)** | SQ | earlyAdolescents | 136 | | | 72 | 0.063 | 0.001 | 0.085 | 0.143 |
| **center (OF)** | SQ | lateAdolescents | 134 | 68 | 25 | | 0.042 | 0.000 | 0.041 | 0.081 |
| **center (OF)** | SQ | adults | 81 | | 22 | | 0.039 | 0.003 | 0.070 | 0.106 |
| **center (OF)*** | HQ | weanlings | 150 | - | - | - | - | - | 0.011 | 0.000 |
| **center (OF)** | HQ | earlyAdolescents | 166 | | | 84 | 0.127 | 0.016 | 0.054 | 0.174 |
| **center (OF)** | HQ | lateAdolescents | 149 | | | 73 | 0.127 | 0.002 | 0.042 | 0.164 |
| **center (OF)*** | HQ | adults | 79 | - | - | - | - | - | 0.019 | 0.000 |
| **brightarm (EP)** | SQ | weanlings | 114 | | | 62 | 0.154 | 0.009 | 0.023 | 0.174 |
| **brightarm (EP)** | SQ | earlyAdolescents | 125 | | 26 | | 0.074 | 0.002 | 0.016 | 0.089 |
| **brightarm (EP)** | SQ | lateAdolescents | 133 | 67 | 25 | | 0.109 | 0.030 | 0.013 | 0.121 |
| **brightarm (EP)** | SQ | adults | 82 | | | 45 | 0.298 | 0.064 | 0.005 | 0.302 |
| **brightarm (EP)** | HQ | weanlings | 150 | 82 | 29 | | 0.129 | 0.038 | 0.007 | 0.134 |
| **brightarm (EP)** | HQ | earlyAdolescents | 163 | 92 | 30 | | 0.265 | 0.084 | 0.028 | 0.286 |
| **brightarm (EP)** | HQ | lateAdolescents | 146 | | | 73 | 0.084 | 0.000 | 0.018 | 0.100 |
| **brightarm (EP)*** | HQ | adults | 78 | - | - | - | - | - | 0.021 | 0.000 |

*Table A18: Meta information about the models. The first column indicates the number of cases per model. Columns two to six indicate the nested mixed effects structure and the number of mice per grouping levels. Columns seven and eight report the adjusted and unadjusted interclass correlation coefficient. The last two columns indicate the marginal and conditional r-squared. The former corresponds to the variance explained by the fixed effects only and the latter by the entire model including both fixed and random effects. For models that were estimated as generalized linear models (GLMs; marked by *) instead of mixed-effects models, variance explained was computed as Mc Faden's pseudo R squared.*

Exploratory analysis testing whether the food switch moderates changes in behavior across development.

Using data from experiments I and II – testing changes from weaning to early adolescence

**Table A19. Regression output (experiments I&II)**

| | | Kenward-Roger Estimates | | | | | Bootstrapped Estimates | | | |
|---|---|---|---|---|---|---|---|---|---|---|
| | **Starting food** | **Estimate** | ***SE*** | **df** | ***t*** | ***p*** | **Estimate** | **95% CI** | ***p*** | ***p adj.*** |
| **distance (OF)** | | | | | | | | | | |
| sexm | SQ | -91.48 | 164.61 | 45.38 | -0.56 | 0.581 | -89.14 | (-420.66, 233.502) | 0.581 | 0.846 |
| switch_time pregnancy | SQ | -70.75 | 381.02 | 41.27 | -0.19 | 0.854 | -77.88 | (-768.328, 664.59) | 0.838 | 0.858 |
| switch_time birth | SQ | -68.23 | 313.25 | 73.97 | -0.22 | 0.828 | -72.82 | (-658.563, 541.018) | 0.812 | 0.812 |
| Testage 9week | SQ | -721.17 | 119.53 | 112.82 | -6.03 | **<0.001** | -724.08 | **(-958.341, -484.07)** | **<0.001** | **<0.001** |
| switch_time pregnancy:testage 9week | SQ | 518.01 | 179.45 | 110.45 | 2.89 | **0.005** | 515.07 | **(164.984, 882.091)** | **0.002** | **0.010** |
| switch_time birth:testage 9week | SQ | 463.72 | 215.29 | 111.55 | 2.15 | **0.033** | 466.95 | **(43.875, 899.94)** | **0.028** | 0.112 |
| sexm | HQ | -105.33 | 137.74 | 69.80 | -0.76 | 0.447 | -104.70 | (-366.51, 158.052) | 0.447 | 0.913 |
| switch_time pregnancy | HQ | -748.21 | 217.45 | 39.46 | -3.44 | **0.001** | -747.21 | **(-1181.435, -336.869)** | **<0.001** | **0.003** |
| switch_time birth | HQ | -541.13 | 268.38 | 74.57 | -2.02 | **0.047** | -541.58 | **(-1047.092, -14.987)** | **0.044** | 0.089 |
| testage 9week | HQ | -419.35 | 168.58 | 155.99 | -2.49 | **0.014** | -420.44 | **(-748.04, -92.12)** | **0.013** | **0.030** |
| switch_time pregnancy:testage 9week | HQ | 144.10 | 236.30 | 147.05 | 0.61 | 0.543 | 145.39 | (-313.807, 603.587) | 0.526 | 0.826 |
| switch_time birth:testage 9week | HQ | -95.21 | 302.50 | 144.64 | -0.31 | 0.753 | -91.85 | (-694.127, 484.099) | 0.760 | 0.914 |
| **center (OF)** | | | | | | | | | | |
| sexm | SQ | 0.00 | 0.14 | 111.37 | 0.02 | 0.987 | 0.00 | (-0.25, 0.27) | 0.981 | 0.981 |
| switch_time pregnancy | SQ | 0.06 | 0.20 | 22.10 | 0.28 | 0.779 | 0.06 | (-0.33, 0.44) | 0.775 | 0.858 |
| switch_time birth | SQ | 0.24 | 0.23 | 116.62 | 1.04 | 0.300 | 0.24 | (-0.2, 0.67) | 0.282 | 0.735 |
| testage 9week | SQ | -0.24 | 0.15 | 117.43 | -1.57 | 0.120 | -0.24 | (-0.52, 0.06) | 0.124 | 0.247 |
| switch_time pregnancy:testage 9week | SQ | 0.36 | 0.23 | 114.21 | 1.58 | 0.118 | 0.36 | (-0.1, 0.81) | 0.121 | 0.242 |
| switch_time birth:testage 9week | SQ | 0.25 | 0.27 | 116.63 | 0.90 | 0.369 | 0.25 | (-0.29, 0.78) | 0.358 | 0.483 |
| sexm | HQ | 0.06 | 0.13 | 74.36 | 0.52 | 0.607 | 0.07 | (-0.18, 0.31) | 0.587 | 0.913 |
| switch_time pregnancy | HQ | 0.16 | 0.17 | 147.50 | 0.97 | 0.336 | 0.16 | (-0.16, 0.5) | 0.342 | 0.455 |
| switch_time birth | HQ | 0.14 | 0.22 | 147.80 | 0.66 | 0.508 | 0.14 | (-0.28, 0.57) | 0.518 | 0.518 |
| testage 9week | HQ | -0.26 | 0.15 | 155.94 | -1.75 | 0.082 | -0.26 | (-0.55, 0.02) | 0.076 | 0.102 |
| switch_time pregnancy:testage 9week | HQ | 0.35 | 0.20 | 146.89 | 1.71 | 0.090 | 0.35 | (-0.05, 0.76) | 0.080 | 0.322 |
| switch_time birth:testage 9week | HQ | -0.03 | 0.26 | 144.25 | -0.11 | 0.913 | -0.03 | (-0.54, 0.49) | 0.914 | 0.914 |
| **distance (EP)** | | | | | | | | | | |
| sexm | SQ | -329.20 | 151.77 | 44.53 | -2.17 | **0.035** | -329.04 | **(-617.032, -31.526)** | **0.030** | 0.122 |
| switch_time pregnancy | SQ | -65.83 | 338.69 | 42.04 | -0.19 | 0.847 | -63.87 | (-703.702, 589.359) | 0.858 | 0.858 |
| switch_time birth | SQ | 186.08 | 304.15 | 98.19 | 0.61 | 0.542 | 180.42 | (-405.817, 773.368) | 0.552 | 0.735 |
| testage 9week | SQ | -38.98 | 189.46 | 104.44 | -0.21 | 0.837 | -38.40 | (-408.105, 346.706) | 0.841 | 0.841 |
| switch_time pregnancy:testage 9week | SQ | 348.81 | 296.33 | 109.56 | 1.18 | 0.242 | 344.33 | (-250.647, 932.203) | 0.252 | 0.337 |
| switch_time birth:testage 9week | SQ | -0.99 | 337.16 | 103.24 | -0.00 | 0.998 | 3.74 | (-661.063, 654.439) | 0.991 | 0.991 |
| sexm | HQ | 16.96 | 144.27 | 67.96 | 0.12 | 0.907 | 16.27 | (-262.681, 295.339) | 0.913 | 0.913 |

**Table A19. Regression output (experiments I&II)**

| | | Kenward-Roger Estimates | | | | | Bootstrapped Estimates | | | |
|---|---|---|---|---|---|---|---|---|---|---|
| | **Starting food** | **Estimate** | ***SE*** | **df** | ***t*** | ***p*** | **Estimate** | **95% CI** | ***p*** | ***p adj.*** |
| switch_time pregnancy | HQ | -321.97 | 249.36 | 39.88 | -1.29 | 0.204 | -321.52 | (-805.867, 173.9) | 0.196 | 0.391 |
| switch_time birth | HQ | -570.91 | 299.97 | 86.10 | -1.90 | 0.060 | -572.13 | **(-1154.106, -19.168)** | **0.044** | 0.089 |
| testage 9week | HQ | 9.74 | 190.33 | 156.35 | 0.05 | 0.959 | 11.08 | (-366.112, 375.391) | 0.952 | 0.952 |
| switch_time pregnancy:testage 9week | HQ | -134.13 | 267.71 | 147.54 | -0.50 | 0.617 | -139.15 | (-644.966, 383.538) | 0.620 | 0.826 |
| switch_time birth:testage 9week | HQ | 318.12 | 340.83 | 143.54 | 0.93 | 0.352 | 320.65 | (-326.161, 998.712) | 0.354 | 0.708 |
| **brightarm (EP)** | | | | | | | | | | |
| sexm | SQ | -0.91 | 1.88 | 53.25 | -0.48 | 0.631 | -0.89 | (-4.578, 2.756) | 0.634 | 0.846 |
| switch_time pregnancy | SQ | -3.81 | 2.81 | 141.70 | -1.36 | 0.177 | -3.81 | (-9.324, 1.758) | 0.178 | 0.710 |
| switch_time birth | SQ | -2.60 | 3.30 | 141.26 | -0.79 | 0.431 | -2.65 | (-9.308, 3.981) | 0.422 | 0.735 |
| testage 9week | SQ | 3.41 | 2.57 | 160.75 | 1.33 | 0.187 | 3.39 | (-1.59, 8.439) | 0.186 | 0.247 |
| switch_time pregnancy:testage 9week | SQ | 1.59 | 4.02 | 161.17 | 0.40 | 0.692 | 1.68 | (-6.225, 9.496) | 0.696 | 0.696 |
| switch_time birth:testage 9week | SQ | 4.16 | 4.59 | 160.37 | 0.91 | 0.366 | 4.19 | (-4.792, 13.193) | 0.362 | 0.483 |
| sexm | HQ | -0.18 | 1.78 | 74.90 | -0.10 | 0.918 | -0.20 | (-3.551, 3.257) | 0.913 | 0.913 |
| switch_time pregnancy | HQ | 0.62 | 2.74 | 46.71 | 0.23 | 0.821 | 0.64 | (-4.671, 6.022) | 0.815 | 0.815 |
| switch_time birth | HQ | 2.41 | 3.44 | 80.09 | 0.70 | 0.486 | 2.51 | (-4.512, 9.151) | 0.463 | 0.518 |
| testage 9week | HQ | 5.89 | 2.41 | 158.02 | 2.45 | **0.016** | 5.90 | **(0.889, 10.497)** | **0.015** | **0.030** |
| switch_time pregnancy:testage 9week | HQ | -0.39 | 3.40 | 148.75 | -0.11 | 0.910 | -0.34 | (-7.046, 6.316) | 0.925 | 0.925 |
| switch_time birth:testage 9week | HQ | -7.09 | 4.34 | 144.57 | -1.63 | 0.105 | -7.13 | (-15.561, 1.493) | 0.097 | 0.387 |

*Table A19: Model summary for all outcome variables measured in weanlings and early adolescents. Each combination of outcome variable and starting food corresponds to one model. All variables were fitted with mixed effects models. We attempted to fit each model with nested random intercepts with mice nested in cages and cages in families. If the model did not converge we simplified the random effects structure (see Table A25). We applied a square root transformation to Center (OF) because the distribution of times spent in the center was skewed towards the right. For all mixed models degrees of freedom were computed with the Kenward-Roger approximation. Additionally, we computed bootstrapped parameters estimates, confidence intervals, and p-values. Per model, we applied a correction for multiple testing (FDR correction) to these bootstrapped p-values across the four outcome variables which use the same set of predictors. Switch time and test age were sum-to-zero coded to avoid multicollinearity resulting from their involvement in an interaction. Sex was dummy coded with females as the reference group. Center (OF) and brightarm (EP) are measured in percent.*

**Table A20. Anova summary (experiments I&II)**

| | **Starting food** | **Sum Sq** | **Mean Sq** | **NumDF** | **DenDF** | ***F*** | ***p*** | ***p adj.*** |
|---|---|---|---|---|---|---|---|---|
| **distance (OF)** | | | | | | | | |
| sex | SQ | 106313.69 | 106314 | 1 | 45.38 | 0.31 | 0.581 | 0.841 |
| switch_time | SQ | 137357.75 | 68679 | 2 | 49.53 | 0.20 | 0.820 | 0.820 |
| testage | SQ | 7476699.91 | 7476700 | 1 | 110.60 | 21.72 | **<0.001** | **<0.001** |
| switch_time:testage | SQ | 3340122.65 | 1670061 | 2 | 110.69 | 4.85 | **0.010** | **0.038** |
| sex | HQ | 474270.96 | 474271 | 1 | 69.80 | 0.58 | 0.447 | 0.918 |
| switch_time | HQ | 12693981.86 | 6346991 | 2 | 25.23 | 7.81 | **0.002** | **0.009** |
| testage | HQ | 9963764.85 | 9963765 | 1 | 143.03 | 12.28 | **<0.001** | **0.002** |
| switch_time:testage | HQ | 600343.92 | 300172 | 2 | 143.89 | 0.37 | 0.691 | 0.691 |
| **center (OF)** | | | | | | | | |
| sex | SQ | 0.00 | 0 | 1 | 111.37 | 0.00 | 0.987 | 0.987 |
| switch_time | SQ | 2.55 | 1 | 2 | 17.94 | 2.17 | 0.143 | 0.571 |
| testage | SQ | 0.06 | 0 | 1 | 115.18 | 0.10 | 0.758 | 0.758 |
| switch_time:testage | SQ | 1.51 | 1 | 2 | 115.09 | 1.30 | 0.275 | 0.551 |
| sex | HQ | 0.16 | 0 | 1 | 74.36 | 0.27 | 0.607 | 0.918 |
| switch_time | HQ | 3.69 | 2 | 2 | 73.50 | 3.02 | 0.055 | 0.110 |

**Table A20. Anova summary (experiments I&II)**

| | Starting food | Sum Sq | Mean Sq | NumDF | DenDF | *F* | *p* | *p adj.* |
|---|---|---|---|---|---|---|---|---|
| testage | HQ | 1.36 | 1 | 1 | 142.68 | 2.24 | 0.137 | 0.183 |
| switch_time:testage | HQ | 2.22 | 1 | 2 | 143.60 | 1.82 | 0.166 | 0.455 |
| **distance (EP)** | | | | | | | | |
| sex | SQ | 4045043.59 | 4045044 | 1 | 44.53 | 4.70 | **0.035** | 0.142 |
| switch_time | SQ | 454051.06 | 227026 | 2 | 42.08 | 0.26 | 0.770 | 0.820 |
| testage | SQ | 275844.89 | 275845 | 1 | 106.09 | 0.32 | 0.572 | 0.758 |
| switch_time:testage | SQ | 1369140.33 | 684570 | 2 | 106.60 | 0.80 | 0.454 | 0.605 |
| sex | HQ | 14257.10 | 14257 | 1 | 67.96 | 0.01 | 0.907 | 0.918 |
| switch_time | HQ | 4762882.84 | 2381441 | 2 | 27.71 | 2.30 | 0.119 | 0.159 |
| testage | HQ | 308759.37 | 308759 | 1 | 142.33 | 0.30 | 0.585 | 0.585 |
| switch_time:testage | HQ | 1827355.33 | 913678 | 2 | 143.47 | 0.89 | 0.415 | 0.553 |
| **brightarm (EP)** | | | | | | | | |
| sex | SQ | 37.83 | 38 | 1 | 53.25 | 0.23 | 0.631 | 0.841 |
| switch_time | SQ | 335.21 | 168 | 2 | 53.36 | 1.04 | 0.362 | 0.723 |
| testage | SQ | 1348.41 | 1348 | 1 | 160.62 | 8.34 | **0.004** | **0.009** |
| switch_time:testage | SQ | 134.00 | 67 | 2 | 160.76 | 0.41 | 0.661 | 0.661 |
| sex | HQ | 1.78 | 2 | 1 | 74.90 | 0.01 | 0.918 | 0.918 |
| switch_time | HQ | 50.69 | 25 | 2 | 23.20 | 0.15 | 0.861 | 0.861 |
| testage | HQ | 711.55 | 712 | 1 | 143.34 | 4.23 | **0.042** | 0.083 |
| switch_time:testage | HQ | 503.89 | 252 | 2 | 144.54 | 1.50 | 0.227 | 0.455 |

*Table A20: Analysis of variance summary with Kenward-Roger degrees of freedom. Each combination of outcome variable and starting food corresponds to one model. Per model, we applied a correction for multiple testing (FDR correction) across the four outcome variables which use the same set of predictors. Star symbols (*) indicate models for which it was not possible to estimate a random effects structure. In those cases we estimated a generalized linear model (GLM). Center (OF) and brightarm (EP) are measured in percent.*

**Table A21. Posthoc pairwise comparisons (experiments I&II)**

| | | | Tukey procedure | | | | | | Unadjusted bootstrapped | |
|---|---|---|---|---|---|---|---|---|---|---|
| | Switch time | Starting food | Estimate | *SE* | df | *t* | 95% CI | *p* | Estimate | 95% CI |
| **distance (OF)** | | | | | | | | | | |
| 5week - 9week | never | SQ | 721.17 | 119.53 | 112.82 | 6.03 | **(484.357,957.986)** | **<0.001** | 727.54 | **(486.916,959.595)** |
| 5week - 9week | pregnancy | SQ | 203.16 | 133.85 | 108.60 | 1.52 | (-62.126,468.452) | 0.132 | 203.05 | (-60.814,470.85) |
| 5week - 9week | birth | SQ | 257.46 | 179.13 | 110.86 | 1.44 | (-97.499,612.41) | 0.153 | 258.72 | (-88.855,615.945) |

*Table A21: Posthoc comparisons of significant interactions. Confidence intervals and p-values have been adjusted for multiple comparisons using the Tukey procedure. The last two columns report bootstrapped estimates and confidence intervals. It is important to note that these bootstrapped confidence intervals are not adjusted for multiple testing. Center (OF) and brightarm (EP) are measured in percent.*

**Table A22. Estimated marginal means and medians (experiments I&II)**

| | | | Marginal means | | | | Bootstrapped marginal medians | |
|---|---|---|---|---|---|---|---|---|
| | Testage | Starting food | Mean | *SE* | df | 95% CI | Median | 95% CI |
| **distance (OF)** | | | | | | | | |
| never | 5week | SQ | 3451.82 | 201.88 | 38.18 | (3043.191,3860.455) | 3458.48 | (3059.291,3860.953) |
| never | 9week | SQ | 2730.65 | 206.65 | 40.86 | (2313.263,3148.041) | 2733.32 | (2339.656,3130.409) |
| pregnancy | 5week | SQ | 3381.08 | 330.04 | 32.50 | (2709.206,4052.949) | 3372.63 | (2763.899,4014.688) |
| pregnancy | 9week | SQ | 3177.91 | 330.07 | 32.50 | (2505.991,3849.838) | 3182.13 | (2548.413,3808.608) |
| birth | 5week | SQ | 3383.59 | 278.24 | 59.51 | (2826.927,3940.257) | 3388.87 | (2857.167,3935.218) |
| birth | 9week | SQ | 3126.14 | 278.17 | 60.98 | (2569.903,3682.37) | 3130.13 | (2581.606,3650.256) |

*Table A22: Estimated marginal means of significant interactions. The last two columns indicate bootstrapped marginal medians and confidence intervals. Center (OF) and brightarm (EP) are measured in percent.*

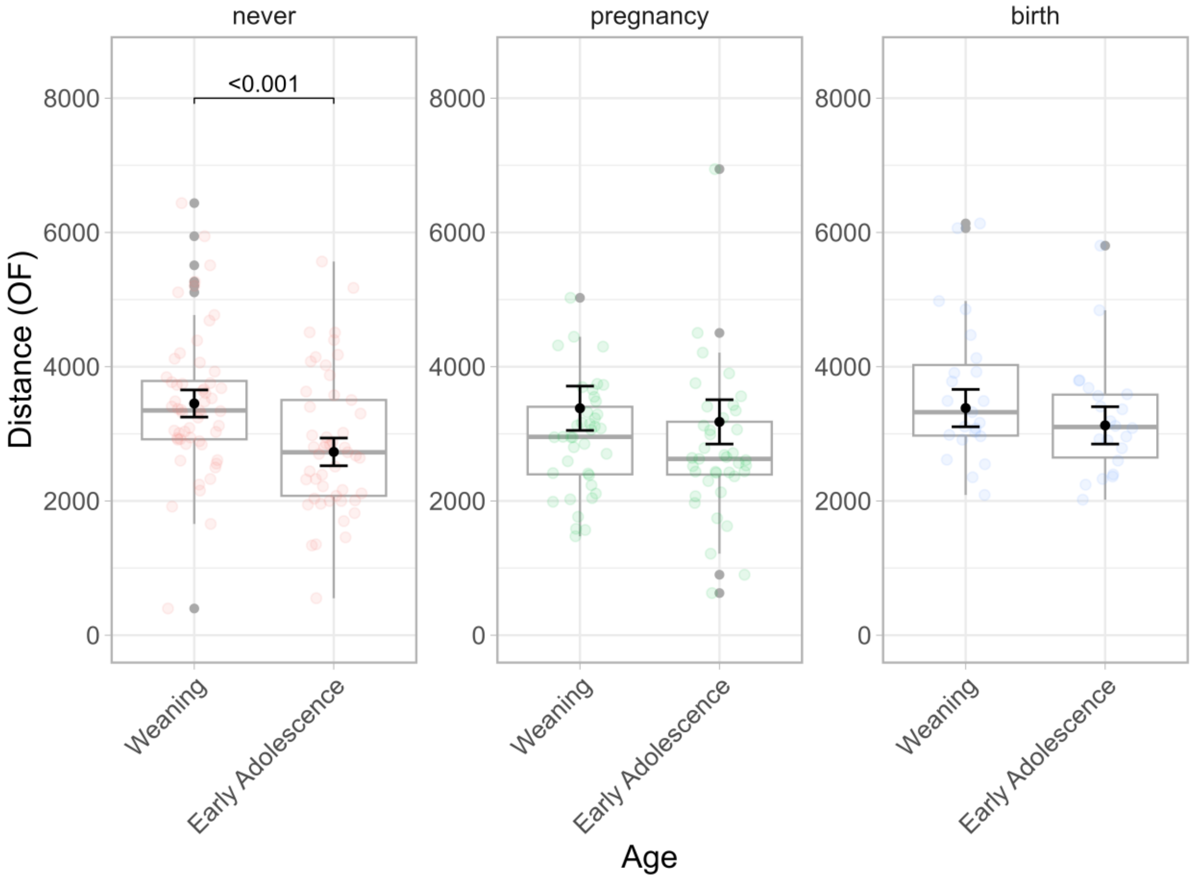


*Figure A4:* Distance covered in the Open Field across development for different switch times in mice that first received standard quality food. Different panels indicate different times during ontogeny at which mice were switched from standard to high quality food, with never corresponding to the control group. Each panel shows boxplots for distance covered in cm (y-axis) across test ages (x-axis). The black points indicate model-based estimated marginal means for each test age and corresponding standard errors. P-values indicate significant differences between test ages. P-values have been corrected for multiple post-hoc comparisons with the Tukey procedure.

#### Using data from experiments I and III - testing changes from early to late adolescence

**Table A23. Regression output (experiments I & III)**

| | | Kenward-Roger Estimates | | | | | Bootstrapped Estimates | | | |
|---|---|---|---|---|---|---|---|---|---|---|
| | **Starting food** | **Estimate** | ***SE*** | **df** | ***t*** | ***p*** | **Estimate** | **95% CI** | ***p*** | ***p adj.*** |
| **distance (OF)** | | | | | | | | | | |
| sexm | SQ | -32.46 | 150.78 | 55.65 | -0.22 | 0.830 | -32.25 | (-315.935, 250.003) | 0.824 | 0.824 |
| switch_time pregnancy | SQ | -222.61 | 339.33 | 24.60 | -0.66 | 0.518 | -227.23 | (-893.377, 445.938) | 0.500 | 0.860 |
| switch_time weanling | SQ | 134.52 | 243.10 | 117.38 | 0.55 | 0.581 | 134.98 | (-343.951, 596.528) | 0.582 | 0.582 |
| testage 13week | SQ | -423.43 | 132.46 | 138.15 | -3.20 | **0.002** | -422.30 | **(-691.901, -168.203)** | **0.001** | **0.005** |
| switch_time pregnancy:testage 13week | SQ | 222.90 | 189.67 | 122.81 | 1.18 | 0.242 | 222.81 | (-155.707, 590.882) | 0.241 | 0.850 |

**Table A23. Regression output (experiments I & III)**

| | | Kenward-Roger Estimates | | | | | Bootstrapped Estimates | | | |
|---|---|---|---|---|---|---|---|---|---|---|
| | Starting food | Estimate | *SE* | df | *t* | *p* | Estimate | 95% CI | *p* | *p adj.* |
| switch_time weanling:testage 13week | SQ | 30.11 | 216.16 | 124.33 | 0.14 | 0.889 | 35.38 | (-395.552, 465.175) | 0.872 | 0.872 |
| sexm | HQ | -205.54 | 144.30 | 71.27 | -1.42 | 0.159 | -201.58 | (-485.289, 69.395) | 0.141 | 0.495 |
| switch_time pregnancy | HQ | -580.54 | 245.14 | 35.25 | -2.37 | **0.023** | -584.52 | **(-1065.031, -103.456)** | **0.016** | **0.029** |
| switch_time weanling | HQ | -110.93 | 262.64 | 145.94 | -0.42 | 0.673 | -106.48 | (-621.684, 393.378) | 0.672 | 0.903 |
| testage 13week | HQ | -531.89 | 158.18 | 174.29 | -3.36 | **<0.001** | -531.95 | **(-843.656, -230.099)** | **<0.001** | **<0.001** |
| switch_time pregnancy:testage 13week | HQ | -97.57 | 216.22 | 151.77 | -0.45 | 0.652 | -103.31 | (-517.629, 328.709) | 0.632 | 0.834 |
| switch_time weanling:testage 13week | HQ | 142.57 | 290.64 | 145.02 | 0.49 | 0.625 | 140.22 | (-406.169, 712.812) | 0.642 | 0.642 |
| **center (OF)** | | | | | | | | | | |
| sexm | SQ | 0.18 | 0.15 | 63.03 | 1.20 | 0.236 | 0.18 | (-0.11, 0.48) | 0.230 | 0.460 |
| switch_time pregnancy | SQ | 0.43 | 0.24 | 25.10 | 1.79 | 0.085 | 0.42 | (0, 0.9) | 0.051 | 0.205 |
| switch_time weanling | SQ | 0.77 | 0.26 | 148.88 | 3.02 | **0.003** | 0.77 | **(0.27, 1.27)** | **0.002** | **0.006** |
| testage 13week | SQ | -0.02 | 0.18 | 161.54 | -0.09 | 0.929 | -0.02 | (-0.37, 0.33) | 0.921 | 0.921 |
| switch_time pregnancy:testage 13week | SQ | 0.06 | 0.27 | 137.88 | 0.23 | 0.817 | 0.06 | (-0.46, 0.58) | 0.817 | 0.850 |
| switch_time weanling:testage 13week | SQ | -0.60 | 0.31 | 138.08 | -1.96 | 0.052 | -0.60 | **(-1.2, -0.01)** | **0.047** | 0.126 |
| sexm | HQ | 0.07 | 0.14 | 74.29 | 0.50 | 0.619 | 0.07 | (-0.2, 0.33) | 0.618 | 0.618 |
| switch_time pregnancy | HQ | 0.50 | 0.19 | 154.69 | 2.62 | **0.010** | 0.50 | **(0.12, 0.86)** | **0.009** | **0.029** |
| switch_time weanling | HQ | 0.03 | 0.25 | 150.70 | 0.11 | 0.910 | 0.03 | (-0.45, 0.52) | 0.903 | 0.903 |
| testage 13week | HQ | 0.09 | 0.17 | 183.74 | 0.56 | 0.576 | 0.09 | (-0.23, 0.41) | 0.564 | 0.564 |
| switch_time pregnancy:testage 13week | HQ | -0.18 | 0.23 | 158.58 | -0.78 | 0.435 | -0.18 | (-0.63, 0.26) | 0.426 | 0.834 |
| switch_time weanling:testage 13week | HQ | -0.26 | 0.31 | 151.97 | -0.83 | 0.407 | -0.26 | (-0.88, 0.34) | 0.398 | 0.642 |
| **distance (EP)** | | | | | | | | | | |
| sexm | SQ | -235.12 | 187.82 | 56.30 | -1.25 | 0.216 | -234.59 | (-604.776, 118.654) | 0.206 | 0.460 |
| switch_time pregnancy | SQ | -153.34 | 381.40 | 25.95 | -0.40 | 0.691 | -160.01 | (-903.897, 587.94) | 0.675 | 0.860 |
| switch_time weanling | SQ | -427.86 | 304.25 | 122.31 | -1.41 | 0.162 | -442.29 | (-1020.484, 170.511) | 0.148 | 0.295 |
| testage 13week | SQ | -424.03 | 176.10 | 126.91 | -2.41 | **0.017** | -428.60 | **(-761.532, -86.211)** | **0.015** | **0.030** |
| switch_time pregnancy:testage 13week | SQ | -52.93 | 265.93 | 117.18 | -0.20 | 0.843 | -49.21 | (-561.724, 485.768) | 0.850 | 0.850 |
| switch_time weanling:testage 13week | SQ | 544.80 | 291.04 | 115.31 | 1.87 | 0.064 | 547.65 | (-23.27, 1118.562) | 0.063 | 0.126 |
| sexm | HQ | -163.63 | 179.15 | 79.64 | -0.91 | 0.364 | -161.98 | (-513.764, 190.89) | 0.372 | 0.495 |
| switch_time pregnancy | HQ | -548.98 | 242.62 | 39.65 | -2.26 | **0.029** | -547.47 | **(-1008.498, -70.369)** | **0.022** | **0.029** |
| switch_time weanling | HQ | -99.62 | 310.19 | 144.68 | -0.32 | 0.749 | -96.19 | (-706.441, 504.535) | 0.752 | 0.903 |
| testage 13week | HQ | -268.69 | 184.56 | 169.56 | -1.46 | 0.147 | -268.07 | (-626.669, 92.677) | 0.147 | 0.294 |
| switch_time pregnancy:testage 13week | HQ | -250.92 | 254.26 | 148.07 | -0.99 | 0.325 | -244.81 | (-756.171, 251.245) | 0.322 | 0.834 |
| switch_time weanling:testage 13week | HQ | 211.86 | 336.40 | 140.02 | 0.63 | 0.530 | 200.27 | (-448.569, 872.186) | 0.529 | 0.642 |
| **brightarm (EP)** | | | | | | | | | | |
| sexm | SQ | 0.61 | 2.43 | 123.78 | 0.25 | 0.801 | 0.63 | (-4.091, 5.236) | 0.806 | 0.824 |
| switch_time pregnancy | SQ | -0.83 | 4.29 | 33.14 | -0.19 | 0.848 | -0.80 | (-9.105, 7.311) | 0.860 | 0.860 |
| switch_time weanling | SQ | -3.63 | 4.28 | 187.30 | -0.85 | 0.397 | -3.58 | (-11.723, 4.692) | 0.383 | 0.510 |
| testage 13week | SQ | 5.14 | 3.09 | 145.61 | 1.66 | 0.098 | 5.11 | (-0.751, 11.042) | 0.090 | 0.120 |
| switch_time pregnancy:testage 13week | SQ | -3.39 | 4.86 | 133.45 | -0.70 | 0.486 | -3.41 | (-12.744, 5.988) | 0.470 | 0.850 |

**Table A23. Regression output (experiments I & III)**

| | | Kenward-Roger Estimates | | | | | Bootstrapped Estimates | | | |
|---|---|---|---|---|---|---|---|---|---|---|
| | **Starting food** | **Estimate** | ***SE*** | **df** | ***t*** | ***p*** | **Estimate** | **95% CI** | ***p*** | ***p adj.*** |
| switch_time weanling:testage 13week | SQ | 1.39 | 5.37 | 129.24 | 0.26 | 0.796 | 1.42 | (-9.128, 11.493) | 0.784 | 0.872 |
| sexm | HQ | -2.09 | 2.00 | 77.22 | -1.04 | 0.300 | -2.09 | (-5.955, 1.82) | 0.288 | 0.495 |
| switch_time pregnancy | HQ | 0.42 | 3.05 | 48.29 | 0.14 | 0.891 | 0.37 | (-5.511, 6.492) | 0.901 | 0.901 |
| switch_time weanling | HQ | -2.02 | 3.75 | 172.16 | -0.54 | 0.591 | -2.02 | (-9.316, 5.064) | 0.600 | 0.903 |
| testage 13week | HQ | -2.05 | 2.53 | 180.97 | -0.81 | 0.418 | -2.07 | (-7.113, 2.919) | 0.406 | 0.541 |
| switch_time pregnancy:testage 13week | HQ | -0.87 | 3.55 | 156.09 | -0.24 | 0.807 | -0.77 | (-7.884, 6.056) | 0.834 | 0.834 |
| switch_time weanling:testage 13week | HQ | 4.77 | 4.73 | 147.62 | 1.01 | 0.315 | 4.79 | (-4.606, 14.044) | 0.309 | 0.642 |

*Table A23: Model summary for all outcome variables measured in early and late adolescents. Each combination of outcome variable and starting food corresponds to one model. All variables were fitted with mixed effects models. We attempted to fit each model with nested random intercepts with mice nested in cages and cages in families. If the model did not converge we simplified the random effects structure (see Table 26). We applied a square root transformation to Center (OF) because the distribution of times spent in the center was skewed towards the right. For all mixed models degrees of freedom were computed with the Kenward-Roger approximation. Additionally, we computed bootstrapped parameters estimates, confidence intervals, and p-values. Per model, we applied a correction for multiple testing (FDR correction) to these bootstrapped p-values across the four outcome variables which use the same set of predictors. Switch time and test age were sum-to-zero coded to avoid multicollinearity resulting from their involvement in an interaction. Sex was dummy coded with females as the reference group. Center (OF) and brightarm (EP) are measured in percent.*

**Table A24. Anova summary (experiments I & III)**

| | **Starting food** | **Sum Sq** | **Mean Sq** | **NumDF** | **DenDF** | ***F*** | ***p*** | ***p adj.*** |
|---|---|---|---|---|---|---|---|---|
| **distance (OF)** | | | | | | | | |
| sex | SQ | 16412.73 | 16413 | 1 | 55.65 | 0.05 | 0.830 | 0.830 |
| switch_time | SQ | 247649.77 | 123825 | 2 | 37.70 | 0.35 | 0.710 | 0.785 |
| testage | SQ | 5518185.91 | 5518186 | 1 | 117.09 | 15.58 | **<0.001** | **<0.001** |
| switch_time:testage | SQ | 543260.00 | 271630 | 2 | 119.74 | 0.77 | 0.467 | 0.622 |
| sex | HQ | 1306792.62 | 1306793 | 1 | 71.27 | 2.03 | 0.159 | 0.485 |
| switch_time | HQ | 5578325.78 | 2789163 | 2 | 38.95 | 4.28 | **0.021** | **0.028** |
| testage | HQ | 14617966.37 | 14617966 | 1 | 141.55 | 22.70 | **<0.001** | **<0.001** |
| switch_time:testage | HQ | 475877.41 | 237939 | 2 | 143.31 | 0.37 | 0.692 | 0.692 |
| **center (OF)** | | | | | | | | |
| sex | SQ | 1.10 | 1 | 1 | 63.03 | 1.43 | 0.236 | 0.471 |
| switch_time | SQ | 6.50 | 3 | 2 | 18.63 | 4.16 | **0.032** | 0.129 |
| testage | SQ | 1.94 | 2 | 1 | 129.74 | 2.53 | 0.114 | 0.114 |
| switch_time:testage | SQ | 3.79 | 2 | 2 | 132.25 | 2.47 | 0.088 | 0.217 |
| sex | HQ | 0.19 | 0 | 1 | 74.29 | 0.25 | 0.619 | 0.619 |
| switch_time | HQ | 7.72 | 4 | 2 | 73.01 | 5.06 | **0.009** | **0.017** |
| testage | HQ | 0.16 | 0 | 1 | 147.79 | 0.21 | 0.646 | 0.675 |
| switch_time:testage | HQ | 0.72 | 0 | 2 | 149.85 | 0.47 | 0.625 | 0.692 |
| **distance (EP)** | | | | | | | | |
| sex | SQ | 991272.15 | 991272 | 1 | 56.30 | 1.57 | 0.216 | 0.471 |
| switch_time | SQ | 312300.10 | 156150 | 2 | 33.24 | 0.24 | 0.785 | 0.785 |
| testage | SQ | 3023860.87 | 3023861 | 1 | 110.20 | 4.78 | **0.031** | 0.060 |
| switch_time:testage | SQ | 2864558.49 | 1432279 | 2 | 113.42 | 2.26 | 0.109 | 0.217 |
| sex | HQ | 715319.03 | 715319 | 1 | 79.64 | 0.83 | 0.364 | 0.485 |
| switch_time | HQ | 10523443.50 | 5261722 | 2 | 30.67 | 6.07 | **0.006** | **0.017** |
| testage | HQ | 4265343.85 | 4265344 | 1 | 136.99 | 4.97 | **0.027** | 0.055 |
| switch_time:testage | HQ | 1903309.17 | 951655 | 2 | 138.96 | 1.11 | 0.333 | 0.692 |
| **brightarm (EP)** | | | | | | | | |
| sex | SQ | 14.91 | 15 | 1 | 123.78 | 0.06 | 0.801 | 0.830 |
| switch_time | SQ | 249.74 | 125 | 2 | 21.37 | 0.53 | 0.598 | 0.785 |

**Table A24. Anova summary (experiments I & III)**

| | Starting food | Sum Sq | Mean Sq | NumDF | DenDF | *F* | *p* | *p adj.* |
|---|---|---|---|---|---|---|---|---|
| testage | SQ | 962.57 | 963 | 1 | 123.89 | 4.11 | **0.045** | 0.060 |
| switch_time:testage | SQ | 183.83 | 92 | 2 | 127.56 | 0.39 | 0.676 | 0.676 |
| sex | HQ | 192.20 | 192 | 1 | 77.22 | 1.09 | 0.300 | 0.485 |
| switch_time | HQ | 3.04 | 2 | 2 | 32.46 | 0.01 | 0.991 | 0.991 |
| testage | HQ | 31.01 | 31 | 1 | 143.99 | 0.18 | 0.675 | 0.675 |
| switch_time:testage | HQ | 260.34 | 130 | 2 | 146.40 | 0.74 | 0.480 | 0.692 |

*Table A24: Analysis of variance summary with Kenward-Roger degrees of freedom. Each combination of outcome variable and starting food corresponds to one model. Per model, we applied a correction for multiple testing (FDR correction) across the four outcome variables which use the same set of predictors. Star symbols (*) indicate models for which it was not possible to estimate a random effects structure. In those cases we estimated a generalized linear model (GLM). Center (OF) and brightarm (EP) are measured in percent.*

**Table A25. Meta information models**

| Outcome | food | experiments | n | fullid_cage_family | cage_family | fullid_cage | fullid_family | family | cage | fullid | ICC_adj | ICC_unadj | r2_m | r2_c |
|---|---|---|---|---|---|---|---|---|---|---|---|---|---|---|
| **distance (OF)** | SQ | exp-I-II | 228 | 119 | 69 | | | 27 | | | 0.706 | 0.539 | 0.060 | 0.724 |
| **distance (OF)** | SQ | exp-I-III | 248 | 140 | 80 | | | 28 | | | 0.644 | 0.379 | 0.040 | 0.658 |
| **distance (OF)** | HQ | exp-I-II | 292 | | | 158 | | 31 | 80 | | 0.231 | 0.063 | 0.123 | 0.326 |
| **distance (OF)** | HQ | exp-I-III | 290 | 165 | 91 | | | 31 | | | 0.410 | 0.184 | 0.142 | 0.494 |
| **distance (EP)** | SQ | exp-I-II | 213 | 118 | 68 | | | 27 | | | 0.271 | 0.113 | 0.034 | 0.296 |
| **distance (EP)** | SQ | exp-I-III | 236 | 138 | 78 | | | 28 | | | 0.569 | 0.399 | 0.038 | 0.585 |
| **distance (EP)** | HQ | exp-I-II | 289 | | | 158 | | 31 | 80 | | 0.178 | 0.039 | 0.035 | 0.206 |
| **distance (EP)** | HQ | exp-I-III | 284 | 164 | 91 | | | 31 | | | 0.405 | 0.171 | 0.095 | 0.462 |
| **center (OF)** | SQ | exp-I-II | 228 | | | | 117 | 27 | | | 0.276 | 0.060 | 0.035 | 0.301 |
| **center (OF)** | SQ | exp-I-III | 248 | 140 | 80 | | | 28 | | | 0.217 | 0.077 | 0.075 | 0.275 |
| **center (OF)** | HQ | exp-I-II | 292 | | | 158 | | | 80 | | 0.242 | 0.103 | 0.043 | 0.274 |
| **center (OF)** | HQ | exp-I-III | 290 | | | 164 | | | 82 | | 0.201 | 0.051 | 0.050 | 0.242 |
| **brightarm (EP)** | SQ | exp-I-II | 213 | | | | | | 63 | | 0.038 | 0.001 | 0.049 | 0.085 |
| **brightarm (EP)** | SQ | exp-I-III | 236 | | | | 138 | 28 | | | 0.179 | 0.010 | 0.029 | 0.203 |
| **brightarm (EP)** | HQ | exp-I-II | 289 | 158 | 85 | | | 31 | | | 0.086 | 0.001 | 0.037 | 0.120 |
| **brightarm (EP)** | HQ | exp-I-III | 284 | 164 | 91 | | | 31 | | | 0.174 | 0.085 | 0.013 | 0.184 |

*Table A25: Meta information about the models. The first column indicates the number of cases per model. Columns two to six indicate the nested mixed effects structure and the number of mice per grouping levels. Columns seven and eight report the adjusted and unadjusted interclass correlation coefficient. The last two columns indicate the marginal and conditional r-squared. The former corresponds to the variance explained by the fixed effects only and the latter by the entire model including both fixed and random effects.*
